# An unsupervised algorithm for host identification in flaviviruses

**DOI:** 10.1101/2020.04.20.050047

**Authors:** Phuoc Truong, Santiago Garcia-Vallve, Pere Puigbò

## Abstract

Early characterization is essential to control the spread of emerging viruses, such as the Zika Virus outbreak in 2014. A major challenge is the identification of potential hosts for novel viruses. We introduce an algorithm to identify the host range of a virus from its raw genome sequence that will be a useful tool to understand host-virus relationships.

## MAIN

Recent viral pandemics have shown that rapid characterization of the virus is essential during the development of an outbreak ^1–4^. Among other factors, host information is essential for surveillance and control of virus spread. However, emerging viruses are fully characterized only after several confirmed cases occur; this is an inefficient method of deterring current and future outbreaks ^5^. Fast and reliable computational biology methods are needed to develop antiviral treatments, to improve medical diagnoses and to efficiently contain viral outbreaks ^6^. Viral genomes are modulated by high mutation rates ^7^ and by selective forces to adapt their codon usage to that of their hosts, especially when the viruses can infect a wide host range, as in flaviviruses ^8^.

In this article, we introduce an unsupervised algorithm to identify putative virus host ranges based on only genome sequence information (figure 1). The proposed methodology has been tested in 94 viruses of genus *Flavivirus* and 16 potential hosts. Several flaviviruses are major human pathogens, with potential host ranges from vertebrates to arthropods ^9^. Flaviviruses are classified by vector type into mosquito-borne (MBFV), tick-borne (TBFV), insect-only (IOFV) and unknown vector (UVFV) ^10^ flaviviruses. In MBFVs, there exists a paraphyletic subgroup of mosquito-specific viruses ^11^, also known as dual-host insect-only flaviviruses (dhIOFVs). Flaviviruses with the same host type tend to be monophyletic and are subject to the same selective pressures as the host; this situation is reflected in their codon usage and dinucleotide composition ^12^. The most widespread and prevalent flaviviruses include Dengue virus (DENV), West Nile virus (WNV), Japanese encephalitis virus (JEV), and Zika virus (ZKV) ^13^.

**Figure 1.**
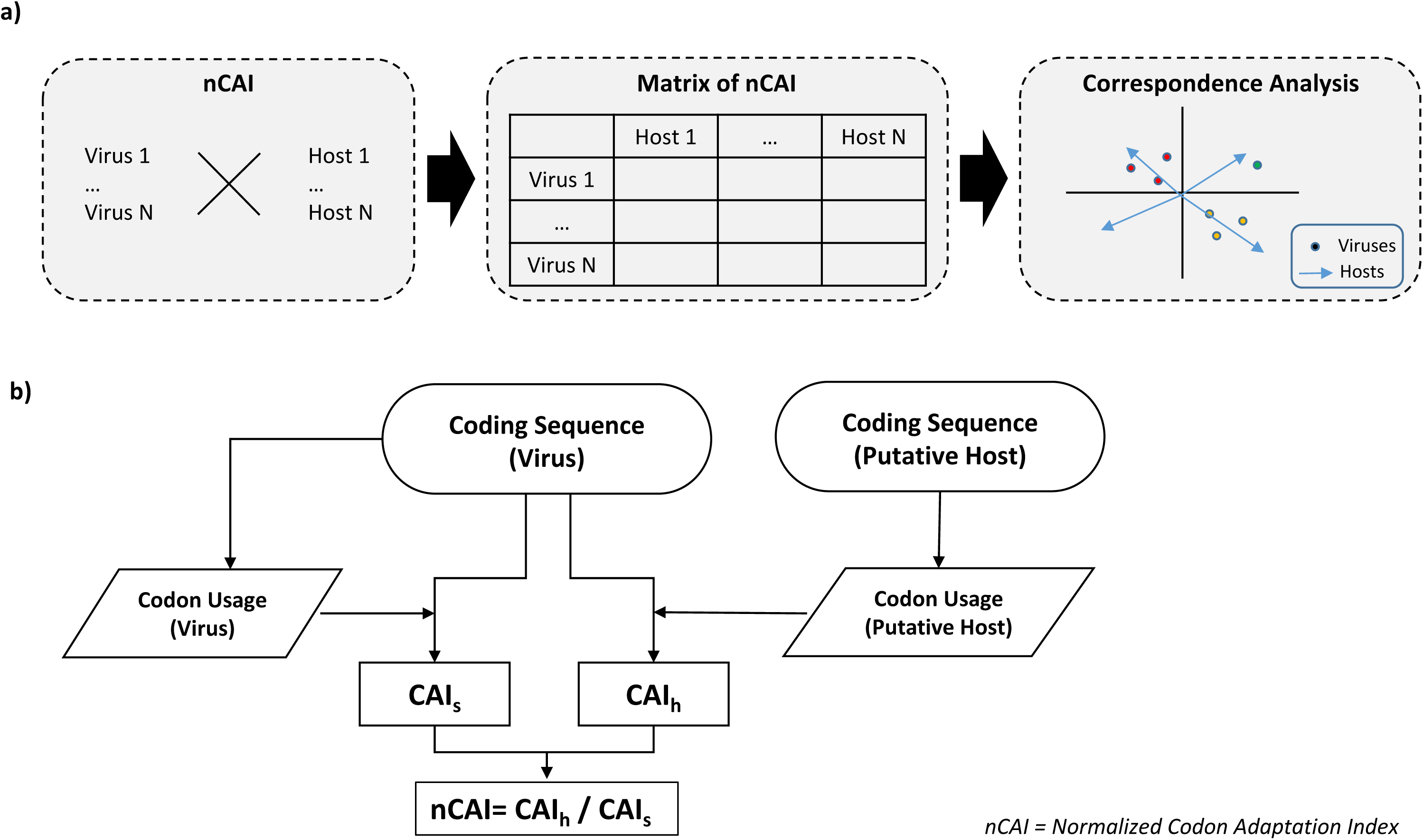
Scheme of the algorithm to calculate the normalized codon adaptation index (nCAI) a) Pipeline to identify putative hosts based on nCAI values. The complete coding sequences of hosts and viruses are used to compute nCAI values, which are put into a table. These values are then subjected to correspondence analysis to identify optimal hosts and, thus, the likelihood of a virus to infect an organism. b) Algorithm to calculate nCAI. The CAI values for possible hosts and viruses of interest are computed from the complete coding sequences (CDSs) and the codon usage tables, which are calculated from the same sequences. The CAI values of the host (CAI_h_) are calculated from virus CDSs and host codon usages, and the CAI values of the viruses (CAI_s_) are computed using virus CDSs and the codon usage values of the viruses themselves. The resulting CAI values are then normalized by dividing each CAI_h_ by its respective CAI_s_.

Several articles suggest that highly similar codon usage frequencies between viruses and hosts are indicative of a high virus-host adaptation level ^14^. Thus, the codon adaptation index (CAI) ^15^ may be a robust indicator to determine putative hosts. Here, we use a normalized CAI (nCAI) to compare codon usage frequencies across virus and host sequences (see methods section). The highest level of optimization is at nCAI=1.0, when the relative use of synonymous codons in the virus and host is identical. Virus-host adaptations are also evaluated with a correspondence analysis (CA) plot (figure 2). Flaviviruses able to infect a wide range of hosts (generalists) tend to be in the center of the plot, whereas host-specific flaviviruses move away from the center towards their optimal hosts (supplementary figures 1–2). Therefore, the nCAI-CA algorithm provides a fast and reliable method of identifying the putative host range of a virus. This method requires only coding sequences (CDSs) without prior knowledge and can be implemented with minimal computational equipment. In addition, we have developed an easy-to-use web server, available at http://ppuigbo.me/programs/CAIcal/nCAI, to calculate nCAI values.

**Figure 2.**
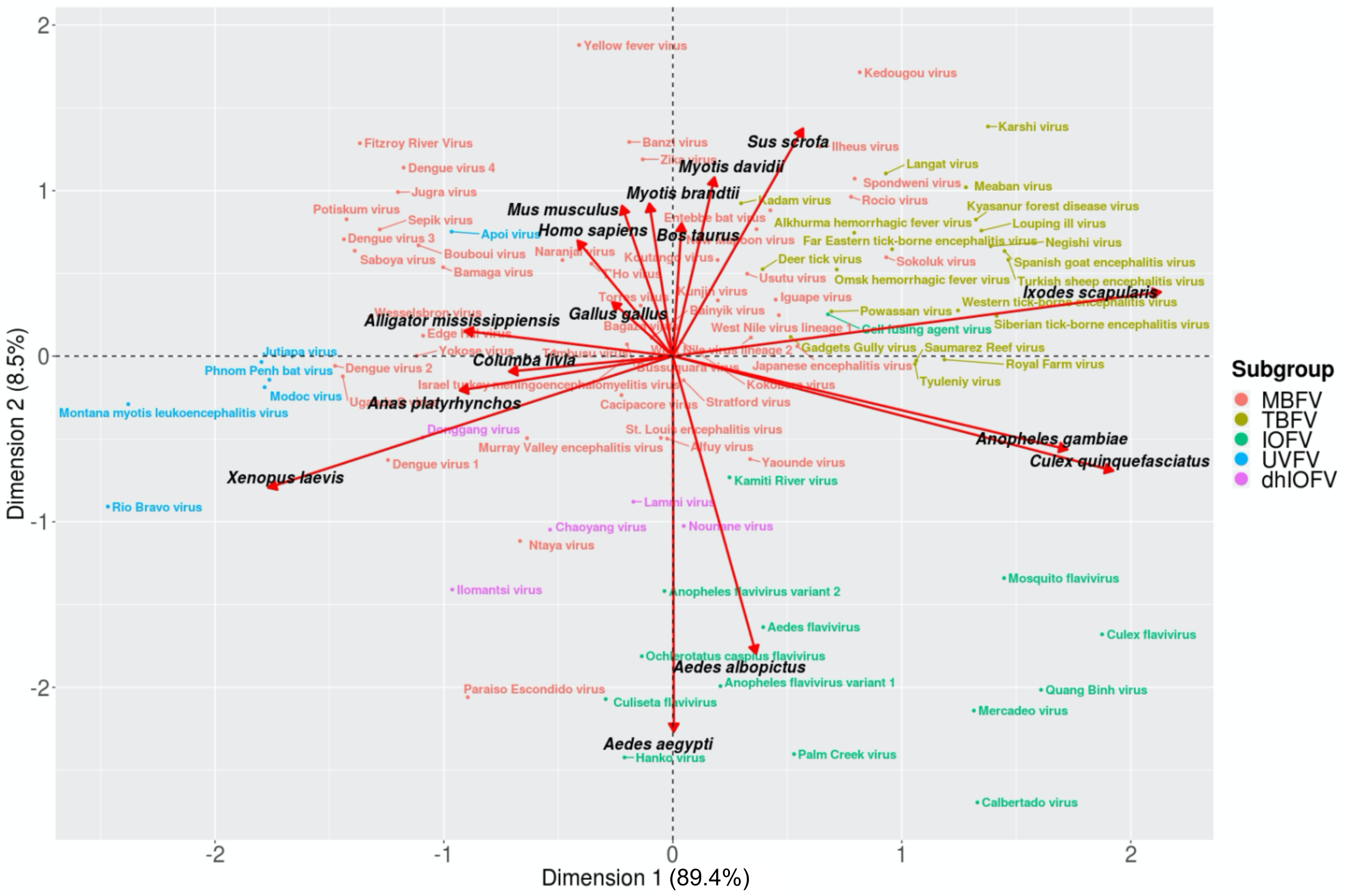
Correspondence analysis of the normalized codon adaptation index (nCAI) values of flaviviruses (genus *Flavivirus*; N = 94). The plot shows that nCAI can differentiate multiple subgroups of flaviviruses based on their degree of codon usage optimization relative to their host organisms. Mosquito-borne flaviviruses are generally optimized for vertebrate hosts, while tick-borne flaviviruses are optimized for ticks, and insect-only flaviviruses are optimized for mosquitoes. Dual-host insect-only flaviviruses show optimization for both mosquitoes and vertebrates, and unknown vector flaviviruses are also optimized for vertebrates. Dimension 1 explains 89.4 percent of the variation, and Dimension 2 explains 8.5 percent of the variation.

First, to assess whether a codon usage methodology could distinguish subgroups within a viral species, we performed nCAI-CA analysis for all the available CDSs of DENV, WNV, JEV, and ZKV, which numbered 4865, 297, 1619 and 494, respectively. Each viral subgroup formed a distinct cluster based on relative synonymous codon usage (supplementary figures 3–4) and GC content (%GC) (supplementary figures 5–7). In addition, we determined the interspecies and intraspecies variability of the relative synonymous codon usage (RSCU) and %GC in the DENV, JEV, WNV and ZKV genomes. The results showed that the RSCU values could differentiate viral subgroups within species and that their distances mostly reflected the evolutionary histories of the viruses (supplementary figure 5). The %GC was not a discriminating factor at the intraspecific level (supplementary figure 6). At the interspecies level, the clustering patterns based on the RSCU were only slightly more similar to the evolutionary histories of the viruses than the %GC (supplementary figure 7).

Next, we used the nCAI-CA algorithm (figure 1) to identify the optimal hosts of 94 flaviviruses based on only complete CDSs and codon usage tables from 16 potential hosts (vertebrates: mammals, birds, reptiles and amphibians; arthropods: mosquitoes and ticks) (supplementary tables 1–3). The nCAI-CA algorithm was able to accurately determine host types for MBFVs and UVFVs (vertebrates), IOFVs (*Aedes* mosquitoes), and TBFVs (*Ixodes scapularis*) (figure 2). The paraphyletic group of dhIOFVs clustered between *Aedes* mosquitoes and vertebrates (supplementary figure 1). The CA plot shows a partial overlap between the MBFV and TBFV groups (supplementary figure 1); however, on average, TBFVs had higher nCAI values (0.813) than MBFVs (0.765) for *I. scapularis*, suggesting a higher degree of optimization for tick hosts (supplementary table 2). The nCAI-CA analysis also revealed unexpected findings for individual viruses, e.g., WNVs clustered within MBFVs but near TBFVs, which aligns with the results of previous infectivity tests ^18^ and some observational studies (supplementary table 1). All the viruses could be classified into two general host groups: vertebrates (MBFVs, TBFVs, UVFVs and dhIOFVs) and mosquitoes (IOFVs) (supplementary figure 2). As expected, no group clustered near *Anopheles gambiae* or *Culex quinquefasciatus*, suggesting that these are not optimal hosts for flaviviruses.

The heat map of nCAI values for all flaviviruses shows common adaptation patterns (supplementary figure 8). Likely optimal hosts (within an nCAI range of 0.9–1.1) include mammals (*Myotis brandtii, M. davidii, Mus musculus, Bos taurus* and *Homo sapiens*) and *Aedes* mosquitoes (*Aedes aegypti* and *A. albopictus*). Unlikely hosts due to low adaptation (nCAI < 0.9) include *C. quinquefasciatus, A. gambiae, I. scapularis* and *Sus scrofa*. These results are in accordance with previous studies and observations, e.g., most MBFVs have a reproductive cycle that includes *Aedes* (host-vector) or *Culex* (vector) mosquitoes and a primary mammalian host (supplementary table 1). Based on these analyses, flaviviruses are potentially less adapted to reproduction in *Culex* mosquitoes due to the differences in %GC between *Culex* and *Aedes* mosquitoes. Moreover, our analysis suggests that TBFV is a group of flaviviruses optimized to reproduce in vertebrates and use ticks as vectors (and we speculate that they may occasionally reproduce in ticks). In general, flavivirus codon usage is overoptimized (nCAI > 1.1) for birds (*Columba livia, Gallus gallus, Anas platyrhynchos*), amphibians (*Xenopus laevis*) and reptiles (*Alligator mississippiensis*).

Despite the accurate results produced by nCAI, there are certain limitations in its application. The algorithm is based on the assumption that there is a selection pressure to optimize the relative use of synonymous codons in the virus. However, it is well known that some viruses use the opposite strategy, and some viruses deoptimize codon usage to hide from host defense mechanisms ^19^. In addition, viruses with overoptimized codon usage (nCAI >> 1) might be explained by multiple factors, e.g., adaptation to multiple hosts, effects of extreme %GC bias or adaptation to highly expressed genes ^20^. Nevertheless, further empirical investigations are necessary to determine reliable confidence intervals for nCAI. Host determination may be uncertain if viruses display approximately equal optimizations for different host types; for example, although dhIOFV codon usage is optimized for both vertebrate and mosquito hosts, they are insect-specific ^11^. Although common host preference patterns are observed, the optimal hosts vary depending on the virus or subgroup and may not reflect documented cases (supplementary table 1). The observed host ranges also do not distinguish between vectors and hosts, and classical phylogenomic methods cannot determine potential hosts without confirmed cases. Our nCAI-based method overcomes this limitation by directly measuring the adaptation of viruses to the translational machinery of their hosts.

In conclusion, this novel algorithm provides a fast and proactive method to assess the potential host ranges and the risk of zoonotic host shift for new and emerging viruses. In flaviviruses, this method distinguishes between arthropod and vertebrate hosts with high accuracy. However, it might produce ambivalent results for viruses undergoing host shifts. Overall, this nCAI-based algorithm may be used as a complement to current phylogenetic methods to monitor current and future outbreaks.

## METHODS

### Normalized codon adaptation index (nCAI)

The optimal host identification algorithm (figure 1b) consists of two phases. In the first phase, the algorithm computes the required codon usage tables through two subroutines: one for the host and the other for the virus. These tables along with complete genomic CDSs are then used as the input data for CAIcal ^21^. This produces CAI data between the virus and host (CAI_h_) using virus CDSs and host codon usage tables and CAI data for the virus itself (CAI_s_) using virus CDSs and virus codon usage tables. In the second phase, the CAI_h_ values are normalized by dividing each by its respective CAI_s_ as follows:

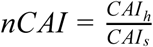

This yields the normalized CAI (nCAI) value, from which the optimal and likely hosts can be inferred depending on how similar the codon usage of a virus is to the codon usage of its host organisms. The nCAI values range between -∞ and +∞, and the optimal value is 1.0, indicating identical codon usage between the virus and host and therefore perfect adaptation to the host. Values above and below 1.0 would indicate over- and underoptimization, respectively, and thus suboptimal adaptation to a host.

### Implementation of the nCAI

The nCAI calculations can be performed with the CAIcal tool in a dedicated web server written in PHP that works on any web browser (http://ppuigbo.me/programs/CAIcal/nCAI). The server requires two sets of inputs: complete DNA or RNA CDSs of the viruses of interest in FASTA format and the codon usage tables of the potential host animals in the format used by the Codon Usage Database ^22^. CAIcal will then output the results in a tab-delimited table with the following values: name of the query sequence (Name), CAI of the virus to a host (CAI_h_), CAI of the virus to itself (CAI_s_), normalized CAI calculated by dividing CAI_h_ by CAI_s_ (nCAI), length of the query sequence (Length), overall %GC, and GC content at the first, second or third nucleotide of each codon (%GC1–3).

### Sequence data

Available flavivirus CDSs and their respective protein sequences were obtained from the RefSeq ^16^ and GenBank ^17^ databases. The viruses were chosen according to phylogenetic studies ^23–25^ and current ICTV classifications ^10^ (supplementary table 1).

### Host and vector organisms

In this study, a vector is defined as an organism capable of transmitting a virus to another type of organism. This definition does not take into account whether the virus is virulent within a vector, i.e., there is no differentiation between a vector and a vector-host. A host is, on the other hand, an organism in which the virus primarily replicates, and it does not directly transmit the virus to another organism of the same type.

The host organisms for this study were chosen based on information primarily provided by the Virus-Host Database ^26^, which includes representative arthropod (mosquitoes and tick) and vertebrate (mammals, birds, reptiles and amphibians) host species. Additionally, a more comprehensive list of hosts and vectors for each flavivirus is included in supplementary table 1. This table includes only confirmed cases of viruses sequenced from an organism or cases in which viruses have successfully infected the cells of a host in a laboratory experiment. It is important to note that not all host animals listed in the database are primary hosts, as they might have acquired the viruses through happenstance.

### Codon usage reference tables

We computed a codon usage reference table for 16 putative hosts representing all possible flavivirus host types among vertebrates (mammals, birds, reptiles and amphibians) and arthropods (mosquitoes and ticks). We analyzed genomes that contained over 10,000 CDSs to reflect actual codon usage frequencies, as well as those of *G. gallus* (6,017) and *S. scrofa* (2,953).

### Genome analysis

The %GC and RSCU values were calculated from the CDSs of the flaviviruses. The RSCU describes the preference bias for a codon to be used to encode an amino acid ^27^. This can be calculated by dividing the observed number of a codon by the expected frequency of the same codon assuming that individual codons for amino acids were used at equal frequency ^28^.

### Correspondence analysis

CA was performed for two different types of nCAI datasets. The first analysis included all known flaviviruses, and the second included separate datasets containing only the values for DENV, JEV, WNV and ZKV. The correspondence analyses were performed with the “ca” package (version 0.70) and then plotted with the “ggplot2” package (version 2.2.1) in R (version 3.4.4).

### Heat maps

The nCAI values of all MBFVs were plotted in a heat map with the “pheatmap” package (version 1.0.10) in R. The virus phylogenetic tree was computed with the following steps: first, the amino acid sequences of each genome were aligned using MUSCLE ^29^. Tree construction was performed with FastTree ^30^. Host trees were built based on NCBI Taxonomy ^31^. Each of the viruses and host organisms were sorted to match their respective phylogenies. Variation test: For the DENV, JEV, WNV and ZKV genomes, their results were clustered based on k-means (5) in the heat map (supplementary figure 9).

### Clustering method

The clustering of each subgroup was performed and visualized by computing centroids based on the multivariate normal distribution of each subgroup with a confidence level of 0.95. This was achieved with the “ggplot2” package (version 2.2.1) in R (version 3.4.4). The virus subgroups included MBFV, TBFV, IOFV, UVFV and dhIOFV, and the host type subgroups were vertebrates, mosquitoes and ticks.

## ACKNOWLEDGMENTS

This work has been supported by funds from the Turku Collegium for Science and Medicine (Turku, Finland).

## AUTHORS CONTRIBUTIONS

Conception and design of the study: PP. Data collection: PT. Data analysis: PP, PT, SGV. Manuscript drafting: PT. Manuscript revision for critical intellectual content: PP, SGV. Writing the final version of the manuscript: PP. All authors read and approved the final manuscript.

