## Supplementary material for "An unsupervised algorithm for host identification in flaviviruses": Supplementary_Material_ppuigbo.pdf

[2] *Present address: Department of Virology, Medicum, Faculty of Medicine, University of Helsinki, Helsinki, Finland.*

### Supplementary table 1. List of flaviviruses (genus *Flavivirus*) and their putative hosts.

| Virus name | Accession codes | | GC3 | Based on the literature | | Correspondence Analysis (CA) | | | |
|---|---|---|---|---|---|---|---|---|---|
| | CDS | AA | | Putative hosts (H) and vectors (V) | Classification | Dim. 1 | Dim. 2 | Centroid classification | The Nearest host |
| Aedes flavivirus | NC_012932.1 | YP_003029843.1 | 0.531 | *Aedes albopictus* (H) [1]<br>*Aedes flavopictus* (H) [1] | IOFV | 0.395 | -1.636 | IOFV / Mosquito | *Aedes albopictus* |
| Alfuy virus | AY898809.1 | AAX82481.1 | 0.525 | *Mus musculus* (H) [1]<br>*Centropus phasianinus* (H) [1]<br>*Mammalia* (H) [2]<br>*Culex pullus* (V) [3]<br>*Culex sitiens* (V) [4] | MBFV | -0.025 | -0.498 | MBFV / Vertebrate, mosquito | *Columba livia* |
| Alkhurma hemorrhagic fever virus | NC_004355.1 | NP_722551.1 | 0.579 | *Ixodes petauristae* (V) [5]<br>*Ixodes ceylonensis* (V) [5]<br>*Homo sapiens* (H) [5, 6] | TBFV | 0.792 | 0.745 | TBFV / Tick, vertebrate | *Sus scrofa* |
| Anopheles flavivirus variant 1 | NC_031327.1 | YP_009305197.1 | 0.525 | *Anopheles gambiae* (H) [1] | IOFV | 0.208 | -1.993 | IOFV / Mosquito | *Aedes albopictus* |
| Anopheles flavivirus variant 2 | KX148547.1 | AOR51360.1 | 0.519 | *Anopheles gambiae* (H) [2] | IOFV | -0.037 | -1.419 | IOFV / Mosquito, vertebrate | *Aedes albopictus* |
| Apoi virus | NC_003676.1 | NP_620045.1 | 0.501 | *Apodemus argenteus* (H) [1] | UVFV | -0.966 | 0.752 | MBFV / Vertebrate | *Homo sapiens* |
| Bagaza virus | NC_012534.1 | YP_002790883.1 | 0.529 | *Culex tritaeniorhynchus* (V) [7]<br>*Homo sapiens* (H) [7]<br>*Alectoris rufa* (H) [8]<br>*Phasianus colchicus* (H) [8] | MBFV | -0.081 | 0.126 | MBFV / Vertebrate, mosquito | *Gallus gallus* |
| Bainyik virus | KM225264.1 | AIJ19433.1 | 0.528 | *Culicidae* (V) [1]<br>*Aedes albopictus* (V) [1]<br>*Mus musculus* (H) [1]<br>*Aedes* sp. (V) [1]<br>Vertebrates (H) [9] | MBFV | -0.093 | 0.111 | MBFV / Vertebrate, mosquito | *Gallus gallus* |
| Bamaga virus | NC_033725.1 | YP_009345036.1 | 0.500 | *Culex sitiens* (V) [1]<br>*Marsupialia* (H) [9] | MBFV | -1.003 | 0.537 | UVFV / Vertebrate | *Alligator mississipiensis* |
| Banzi virus | DQ859056.1 | ABI54472.1 | 0.548 | *Culex rubinotus* (V) [10]<br>*Mansonia africana* (V) [10]<br>*Mesocricetus auratus* (H) [10]<br>*Mastomys natalensis* (H) [10] | MBFV | -0.191 | 1.293 | MBFV / Vertebrate | *Myotis brandtii* |
| Bouboui virus | NC_033693.1 | YP_009344961.1 | 0.493 | *Antilocapra* (H) [1]<br>*Rodentia* (H) [1]<br>*Cercopithecus nictitans* (H) [1]<br>*Papio papio* (H) [1]<br>*Anopheles paludis* (V) [10]<br>*Eretmapodites inornatus* (V) [10]<br>*Aedes* spp. (V) [10]<br>*Culex* spp. (V) [10] | MBFV | -1.111 | 0.669 | MBFV / Vertebrate | *Alligator mississipiensis* |
| Bussuquara virus | NC_009026.2 | YP_001040004.1 | 0.521 | *Chlorocebus aethiops* (H) [1]<br>*Homo sapiens* (H) [1]<br>*Proechimys* spp. (H) [11]<br>*Alouatta belzebul* (H) [12] | MBFV | -0.200 | 0.071 | MBFV / Vertebrate, mosquito | *Gallus gallus* |
| Cacipacore virus | NC_026623.1 | YP_009126874.1 | 0.519 | *Formicarius analis* (H) [1]<br>*Homo sapiens* (H) [13] | MBFV | -0.224 | -0.236 | MBFV / Vertebrate, mosquito | *Columba livia* |

| | | | | | | | | | |
|---|---|---|---|---|---|---|---|---|---|
| Calbertado virus | KX669689.1 | ASA45776.1 | 0.567 | *Culex tarsalis* (H) [2]<br>*Culex pipiens* (H) [14] | IOFV | 1.330 | -2.695 | IOFV / Mosquito | *Aedes albopictus* |
| Cell fusing agent virus | NC_001564.2 | YP_009259257.1 | 0.566 | *Aedes aegypti* (H) [1]<br>*Culicidae* (H) [15] | IOFV | 0.678 | 0.253 | MBFV / Mosquito, vertebrate, tick | *Bos taurus* |
| Chaoyang virus | NC_017086.1 | YP_005454257.1 | 0.492 | *Culicidae* (H) [1] | dhIOFV | -0.536 | -1.048 | dhIOFV / Vertebrate, mosquito | *Anas platyrhynchos* |
| Culex flavivirus | NC_008604.2 | YP_899469.2 | 0.603 | *Culex pipiens* (H) [1] | IOFV | 1.874 | -1.680 | IOFV / Mosquito | *Culex quinquefasciatus* |
| Culiseta flavivirus | NC_030290.1 | YP_009256193.1 | 0.492 | *Culiseta melanura* (H) [1] | IOFV | -0.293 | -2.071 | IOFV / Mosquito | *Aedes aegypti* |
| Deer tick virus | AF311056.1 | AAL32169.1 | 0.561 | *Ixodes scapularis* [1] | TBFV | 0.392 | 0.525 | TBFV / Tick, vertebrate | *Bos taurus* |
| Dengue virus 1 | NC_001477.1 | NP_059433.1 | 0.462 | *Aedes aegypti* (V) [1]<br>*Aedes albopictus* (V) [1]<br>*Homo sapiens* (H) [1] | MBFV | -1.244 | -0.627 | dhIOFV / Vertebrate | *Anas platyrhynchos* |
| Dengue virus 2 | NC_001474.2 | NP_056776.2 | 0.459 | *Aedes aegypti* (V) [1]<br>*Erythrocebus patas* (H) [1]<br>*Homo sapiens* (H) [1]<br>*Aedes furcifer* (V) [1]<br>*Aedes taylori* (V) [1] | MBFV | -1.475 | -0.059 | dhIOFV / Vertebrate | *Anas platyrhynchos* |
| Dengue virus 3 | NC_001475.2 | YP_001621843.1 | 0.468 | *Erythrocebus patas* (H) [1]<br>*Homo sapiens* (H) [1]<br>*Diceromyia* (V) [1]<br>*Aedimorphus* (V) [1]<br>*Stegomyia* (V) [1] | MBFV | -1.437 | 0.706 | MBFV / Vertebrate | *Alligator mississipiensis* |
| Dengue virus 4 | NC_002640.1 | NP_073286.1 | 0.481 | *Aedes aegypti* (V) [1]<br>*Aedes albopictus* (V) [1]<br>*Homo sapiens* (H) [1]<br>*Aedes polynesiensis* (V) [1] | MBFV | -1.176 | 1.138 | MBFV / Vertebrate | *Homo sapiens* |
| Donggang virus | NC_016997.1 | YP_005352889.1 | 0.484 | *Culicidae* (V) [1]<br>*Aedes* sp. (V) [1] | dhIOFV | -1.092 | -0.490 | dhIOFV / Vertebrate | *Anas platyrhynchos* |
| Edge Hill virus | NC_030289.1 | YP_009256192.1 | 0.488 | *Macropodidae* (H) [1]<br>*Culex annulirostris* (V) [10]<br>*Anopheles meraukensis* (V) [10]<br>*Aedes vigilax* (V) [10] | MBFV | -1.091 | 0.123 | UVFV / Vertebrate | *Alligator mississipiensis* |
| Entebbe bat virus | NC_008718.1 | YP_950477.1 | 0.567 | *Chiroptera* (H) [1] | MBFV | 0.426 | 0.881 | MBFV / Vertebrate, tick | *Myotis davidii* |
| Far Eastern tick-borne encephalitis virus | JX498940.1 | AFV41132.1 | 0.589 | *Ixodes persulcatus* (V) [16]<br>*Mus musculus* (H) [16] | TBFV | 0.958 | 0.646 | TBFV / Tick, vertebrate | *Sus scrofa* |
| Fitzroy River Virus | KM361634.1 | AKH03452.1 | 0.483 | *Aedes normanensis* (V) [2]<br>*Anopheles amictus* (V) [17]<br>*Culex annulirostris* (V) [17]<br>*Mammalia* (H) [17]<br>*Aves* (H) [17] | MBFV | -1.367 | 1.286 | MBFV / Vertebrate | *Homo sapiens* |
| Gadgets Gully virus | NC_033723.1 | YP_009345034.1 | 0.553 | *Aves* (H) [1]<br>*Homo sapiens* (H) [1]<br>*Ixodes uriae* (V) [18] | TBFV | 0.515 | 0.116 | TBFV / Tick, vertebrate, mosquito | *Gallus gallus* |
| Hanko virus | NC_030401.1 | YP_009259489.1 | 0.488 | *Culicidae* (H) [1]<br>*Ochlerotatus punctor* (H) [19]<br>*Ochlerotatus caspius* (H) [19] | IOFV | -0.211 | -2.423 | IOFV / Mosquito | *Aedes aegypti* |
| Iguape virus | AY632538.4 | AAV34154.1 | 0.557 | Rodents (H) [20]<br>Sentinel mouse (H) [20] | MBFV | 0.449 | 0.341 | MBFV / Vertebrate, mosquito, tick | *Bos taurus* |

| | | | | | | | | | |
|---|---|---|---|---|---|---|---|---|---|
| | | | | Marsupials (H) [20]<br>Birds (H) [20] | | | | | |
| Ilheus virus | NC_009028.2 | YP_001040006.1 | 0.581 | *Culex* (V) [1]<br>*Haemagogus* (V) [1]<br>*Psorophora* (V) [1]<br>*Aves* (H) [1]<br>*Homo sapiens* (H) [1]<br>*Sabethes* (V) [1]<br>*Ochlerotatus* (V) [1]<br>*Trichoprosopon* (V) [1] | MBFV | 0.643 | 1.266 | MBFV / Vertebrate, tick | *Sus scrofa* |
| Ilomantsi virus | NC_024805.1 | YP_009056847.1 | 0.476 | *Culicidae* (H) [1] | dhIOFV | -0.963 | -1.410 | dhIOFV / Vertebrate, mosquito | *Xenopus laevis* |
| Israel turkey meningoencephalomyelitis virus | KC734549.1 | AGV15505.1 | 0.522 | *Meleagris gallopavo* (H) [2]<br>*Ochlerotatus caspius* (V) [21]<br>*Culicoides imicola* (V) [21]<br>*Culex pipiens* (V) [21]<br>*Phlebotomus papatasi* (V) [21]<br>*Culicoides distinctipennis* (V) [22] | MBFV | -0.218 | -0.028 | MBFV / Vertebrate, mosquito | *Gallus gallus* |
| Japanese encephalitis virus | NC_001437.1 | NP_059434.1 | 0.557 | *Culex tritaeniorhynchus* (V) [1]<br>*Ardeidae* (H) [1]<br>*Homo sapiens* (H) [1]<br>*Equus caballus* (H) [1]<br>*Sus scrofa* (H) [1]<br>*Bos Taurus* (H) [1]<br>*Culex gelidus* (V) [1] | MBFV | 0.544 | 0.060 | MBFV / Vertebrate, mosquito, tick | *Gallus gallus* |
| Jugra virus | NC_033699.1 | YP_009344969.1 | 0.491 | *Cynopterus brachyotis* (H) [1]<br>*Aedes* sp. (V) [10]<br>*Uranotaenia* sp. (V) [10] | MBFV | -1.201 | 0.991 | MBFV / Vertebrate | *Homo sapiens* |
| Jutiapa virus | NC_026620.1 | YP_009126871.1 | 0.447 | *Sigmodon hispidus* (H) [1] | UVFV | -1.797 | -0.035 | UVFV / Vertebrate | *Xenopus laevis* |
| Kadam virus | NC_033724.1 | YP_009345035.1 | 0.560 | *Homo sapiens* [1]<br>*Rhipicephalus pravus* (V) [23]<br>*Rhipicephalus pulchellus* (V) [24]<br>*Amblyomma variegatum* (V) [24]<br>*Hyalomma dromedariil* (V) [25]<br>*Dermacentor variabilis* (V) [26]<br>*Mus musculus* (H) [26] | TBFV | 0.298 | 0.923 | TBFV / Tick, vertebrate | *Myotis davidii* |
| Kamiti River virus | NC_005064.1 | NP_891560.1 | 0.541 | *Aedes* (H) [1] | IOFV | 0.248 | -0.732 | dhIOFV / Mosquito, vertebrate | *Aedes albopictus* |
| Karshi virus | NC_006947.1 | YP_224133.1 | 0.608 | *Homo sapiens* (H) [1]<br>*Rodentia* (H) [1]<br>*Ornithodoros papillipes* (V) [27]<br>*Mus musculus* (H) [27] | TBFV | 1.376 | 1.387 | TBFV / Tick, vertebrate | *Sus scrofa* |
| Kedougou virus | NC_012533.1 | YP_002790882.1 | 0.595 | *Culicidae* (V) [1]<br>*Aedes dalzieli* (V) [28]<br>*Homo sapiens* (H) [29] | MBFV | 0.816 | 1.715 | TBFV / Vertebrate | *Sus scrofa* |
| Kokobera virus | NC_009029.2 | YP_001040007.1 | 0.527 | *Aedes albopictus* (V) [1]<br>*Macropus* (H) [1]<br>*Wallabia* (H) [1]<br>*Homo sapiens* (H) [1]<br>*Culex annulirostris* (V) [1]<br>*Ochlerotatus vigilax* (V) [1]<br>*Ochlerotatus camptorhynchus* (V) [1] | MBFV | 0.036 | 0.089 | MBFV / Vertebrate, mosquito | *Gallus gallus* |

| | | | | | | | | | |
|---|---|---|---|---|---|---|---|---|---|
| | | | | *Culex sitiens* (V) [30] | | | | | |
| Koutango virus | EU082200.2 | ABW76844.2 | 0.549 | *Gerbilliscus kempi* (H) [31]<br>*Rhipicephalus* (V) [31]<br>*Hyalomma* (V) [31]<br>*Ornithodoros* (V) [31]<br>*Aedes aegypti* (V) [32]<br>*Homo sapiens* (H) [33]<br>*Mastomys* (H) [33]<br>*Lemniscomys striatus* (H) [33] | MBFV | 0.196 | 0.581 | MBFV / Vertebrate, tick | *Bos taurus* |
| Kunjin virus | JX276662.1 | AFR66759.1 | 0.545 | *Culex annulirostis* (V) [34]<br>*Aedes tremulus* (V) [35]<br>*Culex australicus* (V) [36]<br>*Culex squamosus* (V) [37]<br>*Aedes vigilax* (V) [38]<br>*Culex quinquefasciatus* (V) [36]<br>*Homo sapiens* (H) [39]<br>*Equus* (H) [40]<br>Sentinel chicken (H) [41]<br>*Nycticorax caledonicus* (H) [42]<br>*Culex pseudovishnui* (V) [43]<br>*Anatidae* sp. [43] | MBFV | 0.197 | 0.337 | MBFV / Vertebrate, mosquito, tick | *Gallus gallus* |
| Kyasanur forest disease virus | AY323490.1 | AAQ91607.1 | 0.603 | *Homo sapiens* (H) [1]<br>*Semnopithecus entellus* (H) [1]<br>*Haemaphysalis spinigera* (V) [44]<br>*Gallus gallus* (H) [45] | TBFV | 1.324 | 0.825 | TBFV / Tick, vertebrate | *Ixodes scapularis* |
| Lammi virus | NC_024806.1 | YP_009056848.1 | 0.518 | *Culicidae* (H) [1] | dhIOFV | -0.173 | -0.879 | dhIOFV / Vertebrate, mosquito | *Columba livia* |
| Langat virus | NC_003690.1 | NP_620108.1 | 0.592 | *Homo sapiens* (H) [1]<br>*Mus* (H) [1]<br>*Ixodes granulatus* (V) [46]<br>*Haemaphysalis Papuana* (V) [47] | TBFV | 0.930 | 1.103 | TBFV / Tick, vertebrate | *Sus scrofa* |
| Louping ill virus | NC_001809.1 | NP_044677.1 | 0.606 | *Homo sapiens* (H) [1]<br>*Canis lupus familiaris* (H) [1]<br>*Equus caballus* (H) [1]<br>*Sus scrofa* (H) [1]<br>*Bos taurus* (H) [1]<br>*Ovis aries* (H) [1]<br>*Ixodes ricinus* (V) [1]<br>*Cervinae* (H) [1] | TBFV | 1.348 | 0.758 | TBFV / Tick, vertebrate | *Ixodes scapularis* |
| Meaban virus | NC_033721.1 | YP_009345031.1 | 0.600 | *Aves* (H) [1]<br>*Homo sapiens* (H) [1]<br>*Ornithodoros maritimus* (V) [48] | TBFV | 1.280 | 1.020 | TBFV / Tick, vertebrate | *Sus scrofa* |
| Mercadeo virus | NC_027819.1 | YP_009164031.1 | 0.573 | *Culex* (H) [1] | IOFV | 1.315 | -2.140 | IOFV / Mosquito | *Aedes albopictus* |
| Modoc virus | NC_003635.1 | NP_619758.1 | 0.447 | *Homo sapiens* (H) [1]<br>*Peromyscus maniculatus* (H) [1] | UVFV | -1.783 | -0.187 | UVFV / Vertebrate | *Xenopus laevis* |
| Montana myotis leukoencephalitis virus | NC_004119.1 | NP_689391.1 | 0.415 | *Myotis lucifugus* (H) [1] | UVFV | -2.379 | -0.290 | UVFV / Vertebrate | *Xenopus laevis* |
| Mosquito flavivirus | NC_021069.1 | YP_007877501.1 | 0.588 | *Culex tritaeniorhynchus* (H) [1] | IOFV | 1.447 | -1.340 | IOFV / Mosquito | *Culex quinquefasciatus* |
| Murray Valley encephalitis virus | NC_000943.1 | NP_051124.1 | 0.493 | *Homo sapiens* (H) [1]<br>*Culex annulirostris* (V) [1] | MBFV | -0.637 | -0.495 | UVFV / Vertebrate, mosquito | *Columba livia* |

| | | | | | | | | | |
|---|---|---|---|---|---|---|---|---|---|
| Naranjal virus | KF917538.1 | AIU94742.1 | 0.516 | Sentinel hamster (H) [2] | MBFV | -0.482 | 0.580 | MBFV / Vertebrate | *Homo sapiens* |
| Negishi virus | KT224355.1 | ALP82435.1 | 0.607 | *Homo sapiens* (H) [49] | TBFV | 1.388 | 0.664 | TBFV / Tick, vertebrate | *Ixodes scapularis* |
| New Mapoon virus | NC_032088.1 | YP_009328360.1 | 0.553 | *Culicidae* (H) [1]<br>*Culex annulirostris* (H) [1] | MBFV | 0.366 | 0.767 | MBFV / Vertebrate, tick | *Bos taurus* |
| Nounane virus | NC_033715.1 | YP_009345019.1 | 0.531 | *Uranotaenia mashonaensis* (H) [2] | dhIOFV | 0.048 | -1.026 | dhIOFV / Vertebrate, mosquito | *Aedes albopictus* |
| Ntaya virus | NC_018705.3 | YP_006846328.2 | 0.489 | *Homo sapiens* (H) [1]<br>*Mus musculus* (H) [1]<br>*Coquillettidia pseudoconopas* (V) [50]<br>*Uranotaenia alboabdominalis* (V) [50]<br>*Culiseta fraseri* (V) [50]<br>*Coquillettidia aurites* (V) [50]<br>*Aedes simpsoni* (V) [50]<br>*Aedes apicoargenteus* (V) [50]<br>*Aedes africanus* (V) [50]<br>*Aedes albomarginatus* (V) [50]<br>*Lutzia tigripes* (V) [50]<br>*Culex poicilipes* (V) [50]<br>*Culex pruina* (V) [50]<br>*Culex moucheti* (V) [50]<br>*Culex* spp. (V) [50] | MBFV | -0.667 | -1.116 | UVFV / Vertebrate, mosquito | *Anas platyrhynchos* |
| Ochlerotatus caspius flavivirus | NC_034242.1 | YP_009352228.1 | 0.499 | *Ochlerotatus caspius* (H) [1]<br>*Aedes albopictus* (H) [51] | IOFV | -0.136 | -1.812 | IOFV / Mosquito | *Aedes aegypti* |
| Omsk hemorrhagic fever virus | NC_005062.1 | NP_878909.1 | 0.577 | *Ixodes* (V) [1]<br>*Homo sapiens* (H) [1]<br>*Ondatra zibethicus* (H) [1]<br>*Dermacentor reticulatus* (V) [1]<br>*Arvicola amphibius* (H) [1] | TBFV | 0.716 | 0.523 | TBFV / Tick, vertebrate | *Bos taurus* |
| Palm Creek virus | NC_033694.1 | YP_009344962.1 | 0.527 | *Coquillettidia xanthogaster* (H) [1] | IOFV | 0.530 | -2.403 | IOFV / Mosquito | *Aedes aegypti* |
| Paraiso Escondido virus | NC_027999.1 | YP_009169331.1 | 0.473 | *Psathyromyia abonnenci* (H) [1] | MBFV | -0.896 | -2.060 | UVFV / Mosquito | *Aedes aegypti* |
| Phnom Penh bat virus | NC_034007.1 | YP_009350101.1 | 0.444 | *Cynopterus brachyotis* (H) [1] | UVFV | -1.762 | -0.143 | UVFV / Vertebrate | *Xenopus laevis* |
| Potiskum virus | NC_029054.2 | YP_009433741.1 | 0.477 | *Culicidae* (V) [1]<br>*Homo sapiens* (H) [1]<br>*Rodentia* (H) [1]<br>*Gallus gallus domesticus* (H) [52] | MBFV | -1.425 | 0.827 | MBFV / Vertebrate | *Alligator mississipiensis* |
| Powassan virus | NC_003687.1 | NP_620099.1 | 0.574 | *Ixodes scapularis* (V) [1]<br>*Homo sapiens* (H) [1]<br>*Marmota monax* (H) [1]<br>*Ixodes spinipalpis* (V) [1]<br>*Dermacentor andersoni* (V) [1]<br>*Ixodes cookei* (V) [1]<br>*Lepus americanus* (H) [1] | TBFV | 0.693 | 0.270 | TBFV / Tick, vertebrate, mosquito | *Bos taurus* |
| Quang Binh virus | NC_012671.1 | YP_002884239.1 | 0.586 | *Culicidae* (H) [1]<br>*Culex tritaeniorhynchus* (H) [1] | IOFV | 1.608 | -2.015 | IOFV / Mosquito | *Aedes albopictus* |

| | | | | | | | | | |
|---|---|---|---|---|---|---|---|---|---|
| Rio Bravo virus | NC_003675.1 | NP_620044.1 | 0.407 | *Homo sapiens* (H) [1]<br>*Eptesicus fuscus* (H) [1]<br>*Tadarida brasiliensis Mexicana* (H) [1]<br>*Molossus ater* (H) [1] | UVFV | -2.468 | -0.908 | UVFV / Vertebrate | *Xenopus laevis* |
| Rocio virus | AY632542.4 | AAV34158.1 | 0.584 | *Homo sapiens* (H) [53, 54]<br>*Mus musculus* (H) [53]<br>*Zonotrichia capensis* (H) [53]<br>*Psorophora ferox* (V) [55, 56]<br>*Aedes scapularis* (V) [55] | MBFV | 0.779 | 0.963 | MBFV / Vertebrate, tick | *Sus scrofa* |
| Royal Farm virus | DQ235149.1 | ABB90673.1 | 0.585 | *Argas hermanni* (V) [57]<br>*Cricetidae* (H) [57] | TBFV | 1.186 | -0.021 | TBFV / Tick, vertebrate, mosquito | *Anopheles gambiae* |
| Saboya virus | NC_033697.1 | YP_009344967.1 | 0.477 | *Mus musculus* (H) [1]<br>*Jaculus jaculus* (H) [1]<br>*Arvicanthis niloticus* (H) [1]<br>*Mastomys sp.* (H) [1]<br>*Gerbilliscus kempi* (H) [1]<br>*Phlebotomus duboscqi* (V) [58, 59]<br>*Sergentomyia inermis* (V) [59]<br>*Sergentomyia squamipleuris* (V) [59]<br>*Sergentomyia adleri* (V) [59]<br>*Sergentomyia clydei* (V) [59]<br>*Sergentomyia antennata* (V) [59]<br>*Sergentomyia buxtoni* (V) [59]<br>*Sergentomyia dubia* (V) [59]<br>*Sergentomyia schwetzi* (V) [59]<br>*Sergentomyia magna* (V) [59] | MBFV | -1.389 | 0.636 | MBFV / Vertebrate | *Alligator mississipiensis* |
| Saumarez Reef virus | NC_033726.1 | YP_009345037.1 | 0.576 | *Aves* (H) [1]<br>*Homo sapiens* (H) [1]<br>*Ornithodoros capensis* (V) [60]<br>*Ixodes eudyptidis* (V) [60] | TBFV | 1.063 | -0.027 | TBFV / Tick, vertebrate, mosquito | *Anopheles gambiae* |
| Sepik virus | NC_008719.1 | YP_950478.1 | 0.483 | *Culicidae* (V) [1]<br>*Ovis aries* (H) [10]<br>*Homo sapiens* (H) [10] | MBFV | -1.280 | 0.765 | MBFV / Vertebrate | *Alligator mississipiensis* |
| Siberian tick-borne encephalitis virus | L40361.3 | AAF82240.2 | 0.607 | *Ixodes persulcatus* (V) [61] | TBFV | 1.414 | 0.244 | TBFV / Tick, vertebrate | *Ixodes scapularis* |
| Sokoluk virus | NC_026624.1 | YP_009126875.1 | 0.588 | *Pipistrellus pipistrellus* (H) [1] | MBFV | 0.932 | 0.597 | TBFV / Vertebrate, tick | *Ixodes scapularis* |
| Spanish goat encephalitis virus | NC_027709.1 | YP_009162613.1 | 0.610 | *Capra hircus* (H) [1] | TBFV | 1.449 | 0.635 | TBFV / Tick, vertebrate | *Sus scrofa* |
| Spondweni virus | NC_029055.1 | YP_009222008.1 | 0.580 | *Aedes circumluteolus* (V) [1]<br>*Mansonia uniformis* (V) [62]<br>*Homo sapiens* (H) [63, 64] | MBFV | 0.794 | 1.072 | MBFV / Vertebrate, tick | *Sus scrofa* |
| St. Louis encephalitis virus | NC_007580.2 | YP_001008348.1 | 0.524 | *Culex quinquefasciatus* (V) [1]<br>*Dromaius novaehollandiae* (H) [1]<br>*Dasypodidae* (H) [1]<br>*Homo sapiens* (H) [1]<br>*Culex nigripalpus* (V) [1]<br>*Passer domesticus* (H) [1] | MBFV | -0.052 | -0.494 | MBFV / Vertebrate, mosquito | *Columba livia* |
| Stratford virus | KM225263.1 | AIJ19432.1 | 0.529 | *Aedes albopictus* (V) [1]<br>*Macropodidae* (H) [1]<br>*Homo sapiens* (H) [1]<br>*Equus caballus* (H) [1] | MBFV | 0.048 | -0.146 | TBFV / Vertebrate, mosquito | *Gallus gallus* |

| | | | | | | | | | |
|---|---|---|---|---|---|---|---|---|---|
| Tembusu virus | NC_015843.2 | YP_004734464.1 | 0.509 | *Anser* sp. (H) [1]<br>*Culex tritaeniorhynchus* (V) [65]<br>*Culex vishnui* (V) [65]<br>*Culex gelidus* (V) [65]<br>*Culex pipiens* (V) [66] | MBFV | -0.518 | 0.030 | UVFV / Vertebrate, mosquito | *Columba livia* |
| T'Ho virus | NC_034151.1 | YP_009351820.1 | 0.527 | *Culex quinquefasciatus* (V) [1]<br>Vertebrates (H) [67] | MBFV | -0.356 | 0.559 | MBFV / Vertebrate | *Homo sapiens* |
| Torres virus | KM225265.1 | AIJ19434.1 | 0.526 | *Culicidae* (V) [1]<br>*Aedes albopictus* (V) [1]<br>*Culex gelidus* (V) [30]<br>*Sus* (H) [30] | MBFV | -0.141 | 0.306 | TBFV / Vertebrate | *Gallus gallus* |
| Turkish sheep encephalitis virus | DQ235151.1 | ABB90675.1 | 0.613 | *Ovis* (H) [68] | TBFV | 1.465 | 0.582 | TBFV / Tick, vertebrate | *Ixodes scapularis* |
| Tyuleniy virus | NC_023424.1 | YP_009001464.1 | 0.579 | *Ixodes uriae* (V) [1] | TBFV | 1.058 | -0.047 | TBFV / Tick, vertebrate, mosquito | *Anopheles gambiae* |
| Uganda S virus | NC_033698.1 | YP_009344968.1 | 0.464 | *Mus musculus* (H) [1]<br>*Saxicola rubetra* (H) [1]<br>*Aedes longipalpis* (V) [69]<br>*Aedes ingrami* (V) [69]<br>*Aedes natronius* (V) [69]<br>*Macaca mulatta* (H) [69] | MBFV | -1.442 | -0.122 | UVFV / Vertebrate | *Anas platyrhynchos* |
| Usutu virus | NC_006551.1 | YP_164264.1 | 0.551 | *Aedes albopictus* (V) [1]<br>*Culex pipiens* (V) [1]<br>*Turdus merula* (H) [1]<br>*Homo sapiens* (H) [1]<br>*Anopheles maculipennis* (V) [1]<br>*Ochlerotatus caspius* (V) [1]<br>*Coquillettidia aurites* (V) [1]<br>*Mansonia Africana* (V) [1]<br>*Culex neavei* (V) [1]<br>*Culex perexiguus* (V) [1] | MBFV | 0.325 | 0.497 | MBFV / Vertebrate, tick | *Bos taurus* |
| Wesselsbron virus | NC_012735.1 | YP_002922020.1 | 0.477 | *Aedes* (V) [1]<br>*Homo sapiens* (H) [1]<br>*Capra hircus* (H) [1]<br>*Ovis aries* (H) [1] | MBFV | -1.324 | 0.237 | MBFV / Vertebrate | *Alligator mississipiensis* |
| West Nile virus lineage 1 | NC_009942.1 | YP_001527877.1 | 0.560 | *Aedes* (V) [1]<br>*Aves* (H) [1]<br>*Homo sapiens* (H) [1]<br>*Amblyomma variegatum* (V) [1]<br>*Hyalomma marginatum* (V) [1]<br>*Rhipicephalus* (V) [1]<br>*Culex* (V) [1]<br>*Mansonia uniformis* (V) [1]<br>*Mimomyia* (V) [1]<br>*Chlorocebus aethiops* (H) [1]<br>*Mesocricetus auratus* (H) [1]<br>*Bubo scandiacus* (H) [1]<br>*Mus* (H) [1]<br>*Corvidae* [70] | MBFV | 0.463 | 0.248 | MBFV / Vertebrate, mosquito, tick | *Bos taurus* |
| West Nile virus lineage 2 | NC_001563.2 | NP_041724.2 | 0.551 | *Homo sapiens* (H) [1] | MBFV | 0.341 | 0.112 | MBFV / Vertebrate, mosquito, tick | *Gallus gallus* |
| Western tick-borne encephalitis virus | NC_001672.1 | NP_043135.1 | 0.595 | *Homo sapiens* (H) [1]<br>*Mus musculus* (H) [1] | TBFV | 1.245 | 0.276 | TBFV / Tick, vertebrate | *Ixodes scapularis* |

| | | | | | | | | | |
|---|---|---|---|---|---|---|---|---|---|
| Yaounde virus | NC_034018.1 | YP_009350103.1 | 0.545 | *Ixodes ricinus* (V) [1]<br>*Ixodes persulcatus* (V) [1]<br>*Culex nebulosus*(V) [1]<br>*Culex telesilla* (V) [71]<br>*Culex quiarti* (V) [71]<br>*Eretmapodites oedipodeios* (V) [71]<br>*Aedes aegypti* (V) [71]<br>*Culex perfuscus* (V) [71]<br>*Culex pruina* (V) [71]<br>*Culex duttoni* (V) [71]<br>*Bycanistes sharpie* (H) [71]<br>*Aves* (H) [72]<br>*Praomys* (H) [71] | MBFV | 0.338 | -0.622 | dhIOFV / Vertebrate, mosquito | *Gallus gallus* |
| Yellow fever virus | NC_002031.1 | NP_041726.1 | 0.537 | *Aedes aegypti* (V) [1]<br>*Aedes simpsoni* (V) [1]<br>*Homo sapiens* (H) [1]<br>*Aedes luteocephalus* (V) [1]<br>*Simiiformes* (H) [1] | MBFV | -0.409 | 1.879 | MBFV / Vertebrate | *Mus musculus* |
| Yokose virus | NC_005039.1 | NP_872627.1 | 0.485 | *Miniopterus fuliginosus* (H) [1]<br>*Culicidae* (V) [73] | MBFV | -1.119 | 0.002 | MBFV / Vertebrate | *Alligator mississippiensis* |
| Zika virus | NC_012532.1 | YP_002790881.1 | 0.541 | *Aedes aegypti* (V) [1]<br>*Aedes albopictus* (V) [1]<br>*Macaca mulatta* (H) [1]<br>*Homo sapiens* (H) [1]<br>*Mus musculus* (H) [1] | MBFV | -0.131 | 1.189 | MBFV / Vertebrate | *Myotis brandtii* |

[1] Information obtained from Virus-Host Database [74].

[2] Information obtained from GenBank [75].

MBFV = mosquito-borne flavivirus

TBFV = tick-borne flavivirus

IOFV = insect-only flavivirus

UVFV = unknown vector flavivirus

dhIOFV = dual-host IOFV

GC3: Proportion of Guanine+Cytosine at the third position of the codon

**Supplementary table 2. Normalized Codon Adaptation Index values of flaviviruses (genus *Flavivirus*) and their putative hosts.**

| **Virus** **Host** | *Aedes aegypti* | *Aedes albopictus* | *Alligator mississippiensis* | *Anas platyrhynchos* | *Anopheles gambiae* | *Bos taurus* | *Columba livia* | *Culex quinquefasciatus* | *Gallus gallus* | *Homo sapiens* | *Ixodes scapularis* | *Mus musculus* | *Myotis brandtii* | *Myotis davidii* | *Sus scrofa* | *Xenopus laevis* |
|---|---|---|---|---|---|---|---|---|---|---|---|---|---|---|---|---|
| Aedes flavivirus | 0,903 | 0,866 | 0,938 | 0,947 | 0,687 | 0,823 | 0,948 | 0,729 | 0,897 | 0,865 | 0,706 | 0,855 | 0,828 | 0,794 | 0,751 | 0,927 |
| Alfuy virus | 0,958 | 0,923 | 1,023 | 1,029 | 0,739 | 0,890 | 1,030 | 0,772 | 0,978 | 0,937 | 0,746 | 0,930 | 0,899 | 0,861 | 0,813 | 1,004 |
| Alkhurma hemorrhagic fever virus | 0,976 | 0,948 | 1,043 | 1,047 | 0,776 | 0,926 | 1,051 | 0,821 | 1,005 | 0,968 | 0,793 | 0,967 | 0,934 | 0,898 | 0,856 | 1,011 |
| Anopheles flavivirus variant 1 | 0,936 | 0,897 | 0,973 | 0,981 | 0,713 | 0,849 | 0,983 | 0,749 | 0,931 | 0,892 | 0,719 | 0,884 | 0,854 | 0,818 | 0,774 | 0,963 |
| Anopheles flavivirus variant 2 | 0,934 | 0,894 | 0,983 | 0,991 | 0,710 | 0,854 | 0,991 | 0,746 | 0,938 | 0,899 | 0,720 | 0,892 | 0,861 | 0,825 | 0,779 | 0,975 |
| Apoi virus | 0,938 | 0,900 | 1,026 | 1,033 | 0,700 | 0,887 | 1,030 | 0,744 | 0,970 | 0,939 | 0,718 | 0,932 | 0,898 | 0,857 | 0,809 | 1,021 |
| Bagaza virus | 1,031 | 0,993 | 1,109 | 1,114 | 0,793 | 0,966 | 1,115 | 0,836 | 1,059 | 1,019 | 0,803 | 1,008 | 0,976 | 0,936 | 0,885 | 1,082 |
| Bainyik virus | 0,961 | 0,928 | 1,035 | 1,040 | 0,743 | 0,901 | 1,042 | 0,779 | 0,987 | 0,949 | 0,750 | 0,942 | 0,912 | 0,874 | 0,826 | 1,013 |
| Bamaga virus | 0,973 | 0,935 | 1,069 | 1,072 | 0,732 | 0,917 | 1,069 | 0,773 | 1,011 | 0,973 | 0,734 | 0,964 | 0,930 | 0,888 | 0,837 | 1,055 |
| Banzi virus | 0,977 | 0,943 | 1,065 | 1,067 | 0,750 | 0,931 | 1,070 | 0,797 | 1,018 | 0,981 | 0,771 | 0,975 | 0,942 | 0,902 | 0,856 | 1,037 |
| Bouboui virus | 0,949 | 0,910 | 1,043 | 1,047 | 0,710 | 0,896 | 1,043 | 0,745 | 0,985 | 0,950 | 0,723 | 0,943 | 0,907 | 0,867 | 0,814 | 1,036 |
| Bussuquara virus | 0,968 | 0,934 | 1,041 | 1,046 | 0,743 | 0,907 | 1,047 | 0,778 | 0,994 | 0,955 | 0,749 | 0,946 | 0,916 | 0,878 | 0,829 | 1,018 |
| Cacipacore virus | 0,987 | 0,949 | 1,058 | 1,065 | 0,753 | 0,919 | 1,066 | 0,791 | 1,009 | 0,970 | 0,766 | 0,961 | 0,929 | 0,888 | 0,839 | 1,039 |
| Calbertado virus | 1,013 | 0,977 | 1,026 | 1,036 | 0,786 | 0,907 | 1,041 | 0,836 | 0,992 | 0,947 | 0,799 | 0,941 | 0,911 | 0,877 | 0,836 | 0,996 |
| Cell fusing agent virus | 0,913 | 0,879 | 0,965 | 0,971 | 0,710 | 0,856 | 0,975 | 0,753 | 0,929 | 0,896 | 0,736 | 0,889 | 0,861 | 0,830 | 0,787 | 0,937 |
| Chaoyang virus | 1,012 | 0,971 | 1,083 | 1,091 | 0,763 | 0,931 | 1,091 | 0,800 | 1,033 | 0,984 | 0,767 | 0,975 | 0,940 | 0,897 | 0,845 | 1,067 |
| Culex flavivirus | 1,030 | 0,999 | 1,050 | 1,061 | 0,820 | 0,939 | 1,067 | 0,876 | 1,024 | 0,975 | 0,844 | 0,971 | 0,943 | 0,911 | 0,871 | 1,009 |
| Culiseta flavivirus | 0,933 | 0,892 | 0,978 | 0,988 | 0,700 | 0,845 | 0,986 | 0,736 | 0,929 | 0,892 | 0,704 | 0,881 | 0,853 | 0,814 | 0,767 | 0,974 |
| Deer tick virus | 0,992 | 0,961 | 1,065 | 1,070 | 0,776 | 0,938 | 1,073 | 0,824 | 1,022 | 0,983 | 0,794 | 0,978 | 0,946 | 0,909 | 0,865 | 1,039 |
| Dengue virus 1 | 0,993 | 0,949 | 1,078 | 1,086 | 0,737 | 0,921 | 1,080 | 0,764 | 1,016 | 0,979 | 0,743 | 0,964 | 0,930 | 0,886 | 0,830 | 1,078 |
| Dengue virus 2 | 1,014 | 0,968 | 1,112 | 1,122 | 0,751 | 0,947 | 1,115 | 0,778 | 1,046 | 1,008 | 0,758 | 0,992 | 0,959 | 0,913 | 0,854 | 1,111 |
| Dengue virus 3 | 0,985 | 0,941 | 1,092 | 1,099 | 0,735 | 0,932 | 1,096 | 0,761 | 1,031 | 0,991 | 0,749 | 0,976 | 0,943 | 0,899 | 0,841 | 1,085 |
| Dengue virus 4 | 0,978 | 0,937 | 1,080 | 1,088 | 0,734 | 0,931 | 1,086 | 0,765 | 1,024 | 0,988 | 0,750 | 0,974 | 0,942 | 0,899 | 0,843 | 1,069 |
| Donggang virus | 0,906 | 0,864 | 0,979 | 0,987 | 0,666 | 0,838 | 0,984 | 0,705 | 0,927 | 0,887 | 0,680 | 0,881 | 0,847 | 0,808 | 0,760 | 0,975 |
| Edge Hill virus | 0,958 | 0,920 | 1,049 | 1,053 | 0,718 | 0,900 | 1,052 | 0,758 | 0,992 | 0,955 | 0,721 | 0,946 | 0,912 | 0,869 | 0,817 | 1,044 |
| Entebbe bat virus | 0,996 | 0,962 | 1,068 | 1,074 | 0,783 | 0,945 | 1,077 | 0,818 | 1,027 | 0,991 | 0,800 | 0,979 | 0,952 | 0,917 | 0,870 | 1,035 |
| Far Eastern tick-borne encephalitis virus | 1,000 | 0,973 | 1,064 | 1,069 | 0,795 | 0,948 | 1,074 | 0,845 | 1,029 | 0,988 | 0,816 | 0,986 | 0,955 | 0,919 | 0,878 | 1,029 |

| | | | | | | | | | | | | | | | | |
|---|---|---|---|---|---|---|---|---|---|---|---|---|---|---|---|---|
| Fitzroy River Virus | 0,964 | 0,922 | 1,068 | 1,074 | 0,716 | 0,919 | 1,072 | 0,754 | 1,008 | 0,976 | 0,732 | 0,964 | 0,931 | 0,887 | 0,834 | 1,060 |
| Gadgets Gully virus | 0,996 | 0,967 | 1,065 | 1,072 | 0,790 | 0,937 | 1,076 | 0,828 | 1,026 | 0,983 | 0,793 | 0,978 | 0,945 | 0,909 | 0,863 | 1,035 |
| Hanko virus | 0,945 | 0,906 | 0,985 | 0,991 | 0,706 | 0,853 | 0,991 | 0,750 | 0,935 | 0,901 | 0,702 | 0,887 | 0,860 | 0,821 | 0,771 | 0,975 |
| Iguape virus | 1,053 | 1,018 | 1,122 | 1,124 | 0,814 | 0,989 | 1,130 | 0,863 | 1,077 | 1,037 | 0,837 | 1,027 | 0,997 | 0,958 | 0,908 | 1,081 |
| Ilheus virus | 0,999 | 0,965 | 1,074 | 1,075 | 0,781 | 0,949 | 1,080 | 0,829 | 1,035 | 0,994 | 0,817 | 0,990 | 0,957 | 0,921 | 0,877 | 1,029 |
| Ilomantsi virus | 0,923 | 0,882 | 0,989 | 0,998 | 0,682 | 0,843 | 0,995 | 0,718 | 0,937 | 0,894 | 0,692 | 0,887 | 0,852 | 0,812 | 0,764 | 0,989 |
| Israel turkey meningoencephalomyelitis virus | 1,015 | 0,977 | 1,092 | 1,097 | 0,776 | 0,949 | 1,099 | 0,818 | 1,041 | 1,000 | 0,787 | 0,992 | 0,959 | 0,917 | 0,867 | 1,069 |
| Japanese encephalitis virus | 0,947 | 0,916 | 1,004 | 1,009 | 0,736 | 0,886 | 1,014 | 0,778 | 0,965 | 0,928 | 0,756 | 0,922 | 0,893 | 0,857 | 0,815 | 0,973 |
| Jugra virus | 0,959 | 0,919 | 1,063 | 1,067 | 0,723 | 0,911 | 1,063 | 0,754 | 1,001 | 0,968 | 0,734 | 0,959 | 0,923 | 0,880 | 0,829 | 1,057 |
| Jutiapa virus | 0,974 | 0,930 | 1,073 | 1,080 | 0,714 | 0,914 | 1,073 | 0,754 | 1,005 | 0,975 | 0,709 | 0,960 | 0,926 | 0,880 | 0,823 | 1,085 |
| Kadam virus | 0,966 | 0,936 | 1,045 | 1,048 | 0,757 | 0,919 | 1,051 | 0,803 | 1,001 | 0,964 | 0,776 | 0,962 | 0,927 | 0,891 | 0,846 | 1,018 |
| Kamiti River virus | 0,917 | 0,880 | 0,960 | 0,969 | 0,693 | 0,845 | 0,970 | 0,740 | 0,918 | 0,888 | 0,719 | 0,883 | 0,851 | 0,816 | 0,775 | 0,948 |
| Karshi virus | 1,005 | 0,981 | 1,074 | 1,074 | 0,820 | 0,964 | 1,082 | 0,865 | 1,043 | 1,004 | 0,835 | 1,001 | 0,970 | 0,938 | 0,898 | 1,025 |
| Kedougou virus | 1,012 | 0,983 | 1,093 | 1,093 | 0,808 | 0,973 | 1,099 | 0,849 | 1,056 | 1,016 | 0,831 | 1,008 | 0,980 | 0,945 | 0,901 | 1,042 |
| Kokobera virus | 0,980 | 0,948 | 1,050 | 1,056 | 0,755 | 0,921 | 1,059 | 0,800 | 1,004 | 0,968 | 0,764 | 0,956 | 0,930 | 0,891 | 0,843 | 1,027 |
| Koutango virus | 1,009 | 0,974 | 1,081 | 1,088 | 0,774 | 0,950 | 1,089 | 0,822 | 1,037 | 0,997 | 0,805 | 0,991 | 0,959 | 0,920 | 0,873 | 1,051 |
| Kunjin virus | 0,994 | 0,960 | 1,065 | 1,071 | 0,770 | 0,934 | 1,073 | 0,813 | 1,022 | 0,982 | 0,786 | 0,975 | 0,943 | 0,905 | 0,859 | 1,038 |
| Kyasanur forest disease virus | 0,990 | 0,964 | 1,050 | 1,054 | 0,797 | 0,941 | 1,061 | 0,848 | 1,019 | 0,979 | 0,823 | 0,979 | 0,947 | 0,913 | 0,872 | 1,010 |
| Lammi virus | 1,012 | 0,972 | 1,080 | 1,087 | 0,773 | 0,933 | 1,087 | 0,807 | 1,031 | 0,984 | 0,782 | 0,976 | 0,941 | 0,901 | 0,849 | 1,057 |
| Langat virus | 1,000 | 0,974 | 1,074 | 1,076 | 0,803 | 0,956 | 1,081 | 0,849 | 1,039 | 0,997 | 0,821 | 0,996 | 0,962 | 0,927 | 0,885 | 1,037 |
| Louping ill virus | 1,000 | 0,974 | 1,062 | 1,068 | 0,810 | 0,949 | 1,074 | 0,856 | 1,032 | 0,988 | 0,832 | 0,987 | 0,956 | 0,922 | 0,883 | 1,021 |
| Meaban virus | 0,956 | 0,934 | 1,021 | 1,024 | 0,776 | 0,914 | 1,030 | 0,825 | 0,991 | 0,952 | 0,790 | 0,950 | 0,920 | 0,887 | 0,849 | 0,979 |
| Mercadeo virus | 0,944 | 0,912 | 0,964 | 0,973 | 0,736 | 0,855 | 0,978 | 0,786 | 0,932 | 0,891 | 0,751 | 0,884 | 0,858 | 0,827 | 0,788 | 0,936 |
| Modoc virus | 0,988 | 0,943 | 1,086 | 1,091 | 0,721 | 0,924 | 1,086 | 0,763 | 1,014 | 0,986 | 0,718 | 0,971 | 0,937 | 0,889 | 0,833 | 1,096 |
| Montana myotis leukoencephalitis virus | 0,974 | 0,927 | 1,079 | 1,086 | 0,696 | 0,908 | 1,077 | 0,737 | 1,004 | 0,974 | 0,694 | 0,959 | 0,923 | 0,873 | 0,814 | 1,099 |
| Mosquito flavivirus | 1,001 | 0,967 | 1,026 | 1,036 | 0,781 | 0,914 | 1,040 | 0,837 | 0,994 | 0,953 | 0,810 | 0,947 | 0,919 | 0,887 | 0,846 | 0,990 |
| Murray Valley encephalitis virus | 0,972 | 0,934 | 1,047 | 1,055 | 0,736 | 0,903 | 1,054 | 0,771 | 0,996 | 0,955 | 0,737 | 0,945 | 0,913 | 0,872 | 0,821 | 1,036 |
| Naranjal virus | 0,974 | 0,938 | 1,059 | 1,063 | 0,744 | 0,919 | 1,064 | 0,779 | 1,008 | 0,972 | 0,755 | 0,963 | 0,929 | 0,889 | 0,839 | 1,039 |
| Negishi virus | 0,995 | 0,970 | 1,056 | 1,061 | 0,807 | 0,944 | 1,067 | 0,854 | 1,026 | 0,983 | 0,827 | 0,981 | 0,951 | 0,917 | 0,877 | 1,014 |

| | | | | | | | | | | | | | | | | |
|---|---|---|---|---|---|---|---|---|---|---|---|---|---|---|---|---|
| New Mapoon virus | 0,921 | 0,889 | 0,986 | 0,991 | 0,714 | 0,871 | 0,995 | 0,755 | 0,946 | 0,913 | 0,744 | 0,907 | 0,877 | 0,843 | 0,801 | 0,960 |
| Nounane virus | 0,992 | 0,953 | 1,051 | 1,058 | 0,760 | 0,910 | 1,058 | 0,792 | 1,004 | 0,960 | 0,770 | 0,952 | 0,919 | 0,881 | 0,832 | 1,026 |
| Ntaya virus | 1,020 | 0,980 | 1,095 | 1,103 | 0,770 | 0,940 | 1,102 | 0,806 | 1,039 | 0,995 | 0,770 | 0,984 | 0,950 | 0,907 | 0,852 | 1,087 |
| Ochlerotatus caspius flavivirus | 0,938 | 0,899 | 0,983 | 0,990 | 0,704 | 0,853 | 0,990 | 0,749 | 0,935 | 0,900 | 0,708 | 0,890 | 0,861 | 0,823 | 0,775 | 0,972 |
| Omsk hemorrhagic fever virus | 1,021 | 0,991 | 1,091 | 1,095 | 0,808 | 0,964 | 1,099 | 0,855 | 1,050 | 1,009 | 0,827 | 1,007 | 0,972 | 0,937 | 0,893 | 1,059 |
| Palm Creek virus | 0,996 | 0,957 | 1,031 | 1,040 | 0,767 | 0,901 | 1,043 | 0,810 | 0,991 | 0,944 | 0,769 | 0,933 | 0,906 | 0,868 | 0,820 | 1,014 |
| Paraiso Escondido virus | 0,996 | 0,956 | 1,064 | 1,073 | 0,745 | 0,905 | 1,069 | 0,781 | 1,008 | 0,960 | 0,734 | 0,950 | 0,915 | 0,872 | 0,820 | 1,065 |
| Phnom Penh bat virus | 0,959 | 0,917 | 1,054 | 1,060 | 0,705 | 0,899 | 1,054 | 0,739 | 0,987 | 0,958 | 0,696 | 0,942 | 0,910 | 0,865 | 0,809 | 1,063 |
| Potiskum virus | 0,987 | 0,944 | 1,090 | 1,095 | 0,726 | 0,934 | 1,091 | 0,773 | 1,026 | 0,993 | 0,742 | 0,983 | 0,947 | 0,902 | 0,847 | 1,086 |
| Powassan virus | 0,996 | 0,966 | 1,062 | 1,066 | 0,786 | 0,938 | 1,070 | 0,834 | 1,022 | 0,982 | 0,802 | 0,978 | 0,946 | 0,911 | 0,867 | 1,032 |
| Quang Binh virus | 0,979 | 0,946 | 0,992 | 1,003 | 0,763 | 0,885 | 1,006 | 0,818 | 0,961 | 0,922 | 0,791 | 0,918 | 0,889 | 0,858 | 0,818 | 0,961 |
| Rio Bravo virus | 0,988 | 0,938 | 1,085 | 1,094 | 0,699 | 0,912 | 1,085 | 0,740 | 1,009 | 0,979 | 0,694 | 0,962 | 0,926 | 0,875 | 0,816 | 1,110 |
| Rocio virus | 1,009 | 0,978 | 1,079 | 1,081 | 0,793 | 0,957 | 1,088 | 0,841 | 1,041 | 1,001 | 0,825 | 0,993 | 0,965 | 0,929 | 0,885 | 1,037 |
| Royal Farm virus | 0,987 | 0,962 | 1,045 | 1,050 | 0,800 | 0,929 | 1,056 | 0,841 | 1,014 | 0,969 | 0,803 | 0,964 | 0,936 | 0,901 | 0,859 | 1,009 |
| Saboya virus | 0,963 | 0,921 | 1,060 | 1,064 | 0,709 | 0,908 | 1,060 | 0,753 | 0,997 | 0,966 | 0,722 | 0,957 | 0,921 | 0,878 | 0,824 | 1,057 |
| Saumarez Reef virus | 0,945 | 0,920 | 1,001 | 1,009 | 0,751 | 0,889 | 1,014 | 0,806 | 0,969 | 0,927 | 0,773 | 0,925 | 0,895 | 0,861 | 0,820 | 0,971 |
| Sepik virus | 0,960 | 0,920 | 1,056 | 1,061 | 0,713 | 0,908 | 1,060 | 0,750 | 0,997 | 0,964 | 0,723 | 0,951 | 0,920 | 0,876 | 0,825 | 1,047 |
| Siberian tick-borne encephalitis virus | 1,014 | 0,990 | 1,068 | 1,072 | 0,815 | 0,955 | 1,079 | 0,866 | 1,035 | 0,995 | 0,836 | 0,992 | 0,962 | 0,928 | 0,887 | 1,026 |
| Sokoluk virus | 0,975 | 0,945 | 1,037 | 1,039 | 0,780 | 0,922 | 1,043 | 0,813 | 1,000 | 0,964 | 0,790 | 0,953 | 0,929 | 0,895 | 0,851 | 0,996 |
| Spanish goat encephalitis virus | 1,010 | 0,987 | 1,071 | 1,075 | 0,821 | 0,959 | 1,082 | 0,868 | 1,042 | 0,997 | 0,841 | 0,995 | 0,965 | 0,932 | 0,892 | 1,027 |
| Spondweni virus | 0,997 | 0,966 | 1,071 | 1,074 | 0,792 | 0,948 | 1,080 | 0,834 | 1,034 | 0,992 | 0,815 | 0,986 | 0,956 | 0,920 | 0,876 | 1,025 |
| St. Louis encephalitis virus | 1,042 | 1,004 | 1,113 | 1,120 | 0,794 | 0,966 | 1,121 | 0,845 | 1,063 | 1,019 | 0,808 | 1,011 | 0,977 | 0,935 | 0,884 | 1,089 |
| Stratford virus | 0,994 | 0,961 | 1,063 | 1,068 | 0,773 | 0,930 | 1,071 | 0,807 | 1,018 | 0,978 | 0,769 | 0,968 | 0,939 | 0,899 | 0,853 | 1,037 |
| Tembusu virus | 0,981 | 0,944 | 1,065 | 1,069 | 0,750 | 0,920 | 1,069 | 0,787 | 1,013 | 0,973 | 0,752 | 0,961 | 0,930 | 0,890 | 0,837 | 1,047 |
| T'Ho virus | 0,991 | 0,954 | 1,076 | 1,080 | 0,761 | 0,936 | 1,080 | 0,795 | 1,025 | 0,987 | 0,773 | 0,978 | 0,945 | 0,905 | 0,854 | 1,055 |
| Torres virus | 0,956 | 0,924 | 1,031 | 1,034 | 0,741 | 0,900 | 1,035 | 0,774 | 0,982 | 0,949 | 0,739 | 0,940 | 0,910 | 0,872 | 0,825 | 1,008 |
| Turkish sheep encephalitis virus | 1,012 | 0,988 | 1,074 | 1,079 | 0,823 | 0,961 | 1,085 | 0,871 | 1,043 | 0,997 | 0,845 | 0,996 | 0,966 | 0,933 | 0,892 | 1,030 |
| Tyuleniy virus | 1,008 | 0,983 | 1,068 | 1,073 | 0,807 | 0,948 | 1,078 | 0,855 | 1,031 | 0,990 | 0,817 | 0,986 | 0,956 | 0,919 | 0,875 | 1,033 |
| Uganda S virus | 0,953 | 0,911 | 1,045 | 1,050 | 0,704 | 0,890 | 1,046 | 0,741 | 0,982 | 0,948 | 0,706 | 0,936 | 0,902 | 0,859 | 0,805 | 1,046 |
| Usutu virus | 0,965 | 0,932 | 1,031 | 1,037 | 0,747 | 0,908 | 1,040 | 0,792 | 0,992 | 0,953 | 0,767 | 0,947 | 0,917 | 0,880 | 0,835 | 1,000 |

| | | | | | | | | | | | | | | | | |
|---|---|---|---|---|---|---|---|---|---|---|---|---|---|---|---|---|
| Wesselsbron virus | 0,977 | 0,934 | 1,068 | 1,076 | 0,719 | 0,916 | 1,074 | 0,761 | 1,009 | 0,973 | 0,733 | 0,963 | 0,926 | 0,884 | 0,831 | 1,067 |
| West Nile virus lineage 1 | 1,006 | 0,973 | 1,071 | 1,076 | 0,781 | 0,944 | 1,079 | 0,829 | 1,031 | 0,990 | 0,801 | 0,984 | 0,952 | 0,915 | 0,869 | 1,038 |
| West Nile virus lineage 2 | 0,999 | 0,964 | 1,063 | 1,070 | 0,767 | 0,934 | 1,072 | 0,819 | 1,022 | 0,979 | 0,792 | 0,974 | 0,942 | 0,904 | 0,859 | 1,031 |
| Western tick-borne encephalitis virus | 0,980 | 0,954 | 1,037 | 1,043 | 0,789 | 0,924 | 1,049 | 0,835 | 1,006 | 0,962 | 0,805 | 0,959 | 0,931 | 0,898 | 0,857 | 0,999 |
| Yaounde virus | 1,043 | 1,007 | 1,103 | 1,108 | 0,803 | 0,965 | 1,111 | 0,849 | 1,058 | 1,013 | 0,817 | 1,005 | 0,973 | 0,934 | 0,886 | 1,072 |
| Yellow fever virus | 0,955 | 0,921 | 1,053 | 1,055 | 0,732 | 0,920 | 1,054 | 0,778 | 1,001 | 0,970 | 0,755 | 0,962 | 0,930 | 0,891 | 0,844 | 1,026 |
| Yokose virus | 0,973 | 0,930 | 1,057 | 1,065 | 0,725 | 0,909 | 1,061 | 0,756 | 0,999 | 0,965 | 0,732 | 0,951 | 0,920 | 0,878 | 0,825 | 1,054 |
| Zika virus | 0,999 | 0,965 | 1,089 | 1,092 | 0,772 | 0,952 | 1,094 | 0,815 | 1,042 | 1,001 | 0,792 | 0,996 | 0,962 | 0,921 | 0,874 | 1,060 |

**Supplementary table 3. Codon usage tables of all putative hosts.**

| | UUU | UCU | UAU | UGU | UUC | UCC | UAC | UGC | UUA | UCA | UAA | UGA | UUG | UCG | UAG | UGG |
|---|---|---|---|---|---|---|---|---|---|---|---|---|---|---|---|---|
| ***Aedes aegypti*** | 105803 | 67316 | 90691 | 70139 | 205867 | 122844 | 160830 | 89653 | 48482 | 75347 | 8409 | 7820 | 162093 | 148951 | 5239 | 82534 |
| ***Aedes albopictus*** | 97145 | 56914 | 79108 | 63514 | 200599 | 123298 | 163078 | 87769 | 38157 | 65567 | 6720 | 5873 | 153673 | 154171 | 4553 | 80333 |
| ***Alligator mississippiensis*** | 285736 | 266501 | 210156 | 170626 | 299463 | 254351 | 242753 | 208569 | 143195 | 222283 | 9468 | 15055 | 237164 | 70524 | 7528 | 204449 |
| ***Anas platyrhynchos*** | 153243 | 135585 | 95599 | 88295 | 149230 | 126705 | 126168 | 110527 | 73660 | 112384 | 1521 | 2599 | 115508 | 37472 | 1122 | 100505 |
| ***Anopheles gambiae*** | 94614 | 32112 | 52725 | 50824 | 165698 | 108606 | 173108 | 91618 | 26775 | 43545 | 7161 | 6391 | 75745 | 182093 | 4436 | 75670 |
| ***Bos taurus*** | 176217 | 159245 | 118803 | 110703 | 230161 | 202850 | 170291 | 140579 | 79353 | 126638 | 8272 | 17113 | 136301 | 59386 | 6476 | 137993 |
| ***Columba livia*** | 112853 | 92944 | 74363 | 65581 | 113012 | 93945 | 97512 | 76376 | 50686 | 75022 | 3715 | 4794 | 84780 | 28616 | 2286 | 73187 |
| ***Culex quinquefasciatus*** | 123786 | 42919 | 50760 | 60923 | 198511 | 134088 | 195233 | 107214 | 25108 | 52078 | 7306 | 6787 | 134792 | 188984 | 4659 | 86930 |
| ***Gallus gallus*** | 45768 | 38296 | 32211 | 23851 | 54936 | 42683 | 48342 | 36075 | 19129 | 31442 | 2046 | 2986 | 34146 | 14079 | 1281 | 32616 |
| ***Homo sapiens*** | 189379 | 171196 | 132715 | 117458 | 216388 | 196012 | 161579 | 135489 | 86682 | 139095 | 5404 | 9518 | 143507 | 50249 | 4328 | 134648 |
| ***Ixodes scapularis*** | 74374 | 55722 | 31434 | 33921 | 155979 | 105861 | 129415 | 100447 | 19307 | 37110 | 4241 | 8274 | 66146 | 96803 | 4231 | 72965 |
| ***Mus musculus*** | 422153 | 398250 | 298518 | 279729 | 535439 | 444041 | 394074 | 301384 | 165150 | 289799 | 23403 | 40148 | 329668 | 103815 | 19126 | 306619 |
| ***Myotis brandtii*** | 149256 | 134443 | 103108 | 91596 | 189819 | 173931 | 143371 | 113452 | 69465 | 105333 | 5078 | 9823 | 118397 | 45721 | 4583 | 113559 |
| ***Myotis davidii*** | 113232 | 103442 | 76640 | 70839 | 155462 | 144783 | 117529 | 95128 | 51237 | 81258 | 3784 | 8156 | 90795 | 39444 | 3674 | 92755 |
| ***Sus scrofa*** | 18160 | 14246 | 12717 | 10902 | 27973 | 21586 | 22023 | 16776 | 6442 | 10448 | 883 | 1739 | 13518 | 5607 | 628 | 17440 |
| ***Xenopus laevis*** | 475411 | 435671 | 348838 | 278073 | 343215 | 321198 | 290496 | 239628 | 259966 | 343805 | 19075 | 17534 | 332659 | 77483 | 9401 | 245817 |

| | CUU | CCU | CAU | CGU | CUC | CCC | CAC | CGC | CUA | CCA | CAA | CGA | CUG | CCG | CAG | CGG |
|---|---|---|---|---|---|---|---|---|---|---|---|---|---|---|---|---|
| ***Aedes aegypti*** | 81073 | 70190 | 92779 | 70386 | 92845 | 78473 | 111503 | 74523 | 69643 | 116049 | 154187 | 94417 | 248006 | 143421 | 198592 | 92034 |
| ***Aedes albopictus*** | 73743 | 63435 | 85963 | 66043 | 91059 | 79507 | 115067 | 74949 | 67101 | 105136 | 142086 | 89873 | 262230 | 158240 | 205119 | 103979 |
| ***Alligator mississippiensis*** | 234983 | 286349 | 199397 | 80956 | 261744 | 253141 | 224817 | 142817 | 134363 | 313100 | 247331 | 88040 | 592915 | 94622 | 565858 | 148393 |
| ***Anas platyrhynchos*** | 122813 | 143338 | 94079 | 42336 | 135241 | 121613 | 119654 | 60556 | 60117 | 145719 | 122456 | 45366 | 278261 | 46915 | 271533 | 61272 |
| ***Anopheles gambiae*** | 51540 | 33672 | 71298 | 60653 | 110524 | 79021 | 127400 | 144786 | 51475 | 75633 | 91709 | 56430 | 341455 | 213374 | 255693 | 126863 |
| ***Bos taurus*** | 137259 | 183974 | 106989 | 47991 | 226797 | 238009 | 174809 | 122283 | 69555 | 174268 | 124199 | 67416 | 452888 | 98259 | 381244 | 137128 |
| ***Columba livia*** | 83984 | 95097 | 66456 | 29591 | 101401 | 92973 | 87675 | 45990 | 38718 | 99227 | 82943 | 32668 | 206622 | 37795 | 189692 | 48921 |
| ***Culex quinquefasciatus*** | 64004 | 42642 | 58083 | 58474 | 126302 | 97230 | 150387 | 115735 | 44253 | 87713 | 119463 | 75642 | 342737 | 212800 | 255462 | 143812 |
| ***Gallus gallus*** | 33708 | 41672 | 25885 | 14682 | 45753 | 46097 | 39081 | 28305 | 16211 | 42767 | 33018 | 14339 | 104699 | 21091 | 88743 | 26453 |
| ***Homo sapiens*** | 147569 | 198345 | 123609 | 49921 | 212802 | 225420 | 168062 | 115976 | 79488 | 192119 | 140427 | 68859 | 437308 | 81354 | 387120 | 129331 |
| ***Ixodes scapularis*** | 66042 | 55208 | 36505 | 40240 | 159231 | 120308 | 116979 | 101295 | 31749 | 59161 | 59065 | 44503 | 233226 | 106684 | 169406 | 86673 |
| ***Mus musculus*** | 329757 | 450637 | 260637 | 114854 | 495018 | 446868 | 375626 | 229758 | 198032 | 423707 | 293318 | 161412 | 969515 | 151521 | 836320 | 250836 |
| ***Myotis brandtii*** | 114062 | 157094 | 94505 | 38894 | 185980 | 195453 | 149357 | 88661 | 60061 | 148475 | 111991 | 56543 | 375305 | 64984 | 332354 | 107579 |
| ***Myotis davidii*** | 86828 | 124280 | 72079 | 30930 | 154989 | 167094 | 125963 | 77771 | 45385 | 116015 | 84072 | 44250 | 317889 | 60214 | 271506 | 94037 |
| ***Sus scrofa*** | 13109 | 18561 | 9900 | 4792 | 27053 | 25796 | 18267 | 14145 | 6653 | 16692 | 11567 | 6519 | 53901 | 9860 | 40912 | 13943 |
| ***Xenopus laevis*** | 378042 | 377145 | 298817 | 119961 | 274223 | 244929 | 267467 | 120706 | 213125 | 432029 | 395345 | 122409 | 540897 | 91856 | 585594 | 119667 |

| | AUU | ACU | AAU | AGU | AUC | ACC | AAC | AGC | AUA | ACA | AAA | AGA | AUG | ACG | AAG | AGG |
|---|---|---|---|---|---|---|---|---|---|---|---|---|---|---|---|---|
| ***Aedes aegypti*** | 152166 | 86616 | 166027 | 105612 | 207964 | 148111 | 221033 | 123334 | 76072 | 84025 | 220087 | 53146 | 189682 | 133677 | 268281 | 43987 |
| ***Aedes albopictus*** | 137495 | 76150 | 147062 | 100974 | 208043 | 153503 | 225036 | 125677 | 67644 | 75402 | 206848 | 47629 | 186399 | 137826 | 273900 | 44683 |
| ***Alligator mississippiensis*** | 289540 | 245848 | 311827 | 221332 | 318302 | 263582 | 322932 | 326064 | 160920 | 294026 | 477989 | 230179 | 377994 | 97321 | 532404 | 200385 |
| ***Anas platyrhynchos*** | 141066 | 117055 | 148135 | 106071 | 153436 | 121061 | 168842 | 167445 | 80265 | 148079 | 247562 | 118152 | 173849 | 57102 | 251100 | 107972 |
| ***Anopheles gambiae*** | 92725 | 39481 | 100091 | 64028 | 202044 | 140594 | 219692 | 156991 | 61791 | 59333 | 126145 | 23006 | 166014 | 201293 | 262509 | 21939 |
| ***Bos taurus*** | 159826 | 133607 | 163964 | 129267 | 234653 | 210799 | 212732 | 224422 | 76928 | 150192 | 251908 | 130360 | 233655 | 79271 | 350736 | 134871 |
| ***Columba livia*** | 104393 | 80840 | 107072 | 73499 | 121220 | 92966 | 128058 | 113525 | 58143 | 104091 | 182096 | 84864 | 131436 | 42191 | 189730 | 74351 |
| ***Culex quinquefasciatus*** | 128311 | 53265 | 99723 | 87381 | 246469 | 173860 | 278300 | 147234 | 39912 | 52962 | 149090 | 36161 | 187954 | 191512 | 336380 | 40696 |
| ***Gallus gallus*** | 45653 | 36078 | 46039 | 30390 | 59906 | 44951 | 61099 | 54867 | 23805 | 43884 | 74256 | 33289 | 62972 | 20943 | 93393 | 31945 |
| ***Homo sapiens*** | 175259 | 147134 | 190114 | 139465 | 220634 | 203602 | 206688 | 220505 | 82922 | 167136 | 278169 | 133268 | 236510 | 66200 | 353825 | 131616 |
| ***Ixodes scapularis*** | 57705 | 45993 | 45690 | 42553 | 134295 | 111797 | 147940 | 121031 | 31509 | 54105 | 78016 | 39079 | 127674 | 107148 | 208833 | 78311 |
| ***Mus musculus*** | 377698 | 335039 | 382284 | 311331 | 552184 | 465115 | 499149 | 483013 | 180467 | 391437 | 537723 | 297135 | 559953 | 138180 | 825270 | 299472 |
| ***Myotis brandtii*** | 139814 | 114452 | 148430 | 113992 | 199193 | 180990 | 185755 | 185631 | 66531 | 131463 | 226768 | 109231 | 204831 | 65277 | 313098 | 118086 |
| ***Myotis davidii*** | 105319 | 86591 | 110660 | 87677 | 163717 | 149818 | 150586 | 156457 | 49298 | 100435 | 169437 | 82710 | 162715 | 57159 | 249402 | 96344 |
| ***Sus scrofa*** | 15660 | 13038 | 16598 | 11082 | 28754 | 26456 | 25606 | 23327 | 7191 | 14403 | 23681 | 12052 | 25610 | 9022 | 38617 | 13258 |
| ***Xenopus laevis*** | 470669 | 372157 | 520334 | 336460 | 352434 | 296177 | 419539 | 337006 | 300000 | 463817 | 720130 | 342826 | 502530 | 95128 | 601326 | 252104 |

| | GUU | GCU | GAU | GGU | GUC | GCC | GAC | GGC | GUA | GCA | GAA | GGA | GUG | GCG | GAG | GGG |
|---|---|---|---|---|---|---|---|---|---|---|---|---|---|---|---|---|
| ***Aedes aegypti*** | 128092 | 123462 | 238605 | 116851 | 115996 | 177203 | 192964 | 115501 | 75096 | 112842 | 295762 | 166203 | 173024 | 103277 | 220013 | 58749 |
| ***Aedes albopictus*** | 120907 | 114500 | 230285 | 116327 | 117235 | 184026 | 202864 | 117142 | 71956 | 106383 | 283652 | 156762 | 180844 | 114481 | 230877 | 62953 |
| ***Alligator mississippiensis*** | 220062 | 336797 | 425108 | 182259 | 214803 | 357261 | 392225 | 290968 | 146333 | 328225 | 560811 | 297308 | 410333 | 94451 | 643968 | 250922 |
| ***Anas platyrhynchos*** | 117470 | 181596 | 201191 | 94731 | 105498 | 153253 | 180425 | 133781 | 74738 | 173629 | 275108 | 151883 | 198074 | 45370 | 284768 | 115330 |
| ***Anopheles gambiae*** | 64418 | 75543 | 173980 | 125515 | 109118 | 190816 | 212737 | 204626 | 60279 | 106001 | 179852 | 87574 | 229677 | 195713 | 283970 | 78940 |
| ***Bos taurus*** | 113850 | 193365 | 220484 | 111861 | 168797 | 326809 | 288879 | 256275 | 70781 | 160241 | 302609 | 174863 | 312161 | 100065 | 441648 | 185928 |
| ***Columba livia*** | 84579 | 119203 | 148911 | 69163 | 84576 | 113404 | 141019 | 95801 | 50144 | 110554 | 201781 | 109153 | 151071 | 39420 | 213638 | 86177 |
| ***Culex quinquefasciatus*** | 113170 | 86702 | 170362 | 108252 | 153568 | 225445 | 270497 | 167341 | 47010 | 81107 | 221992 | 143559 | 218927 | 184999 | 313182 | 89461 |
| ***Gallus gallus*** | 35593 | 56528 | 68683 | 30898 | 36917 | 62202 | 67783 | 53631 | 21277 | 51713 | 84178 | 47765 | 76624 | 24768 | 111123 | 43513 |
| ***Homo sapiens*** | 121302 | 204091 | 246943 | 117456 | 155761 | 311996 | 278549 | 247607 | 78882 | 178106 | 336665 | 183190 | 305878 | 84501 | 451726 | 182999 |
| ***Ixodes scapularis*** | 56833 | 73718 | 66403 | 60167 | 135465 | 195993 | 232841 | 161342 | 31338 | 81570 | 121651 | 88146 | 190498 | 115433 | 225468 | 82020 |
| ***Mus musculus*** | 262535 | 491093 | 515049 | 280522 | 377902 | 637878 | 638504 | 520069 | 182733 | 388723 | 661498 | 411344 | 696158 | 157124 | 965963 | 372099 |
| ***Myotis brandtii*** | 97606 | 165822 | 196915 | 92699 | 140631 | 262234 | 251134 | 199899 | 60268 | 140268 | 275392 | 144558 | 268064 | 65575 | 386908 | 155884 |
| ***Myotis davidii*** | 74166 | 130534 | 149620 | 72558 | 116940 | 225972 | 209517 | 171189 | 45056 | 110134 | 205844 | 112003 | 222467 | 60004 | 317761 | 132602 |
| ***Sus scrofa*** | 10755 | 19642 | 22309 | 11651 | 20245 | 36983 | 33278 | 29987 | 6447 | 15141 | 27310 | 18686 | 38572 | 10339 | 48046 | 21555 |
| ***Xenopus laevis*** | 351271 | 411579 | 613288 | 253098 | 244083 | 319491 | 432347 | 265240 | 240447 | 424584 | 772321 | 441213 | 434149 | 85444 | 662177 | 270557 |

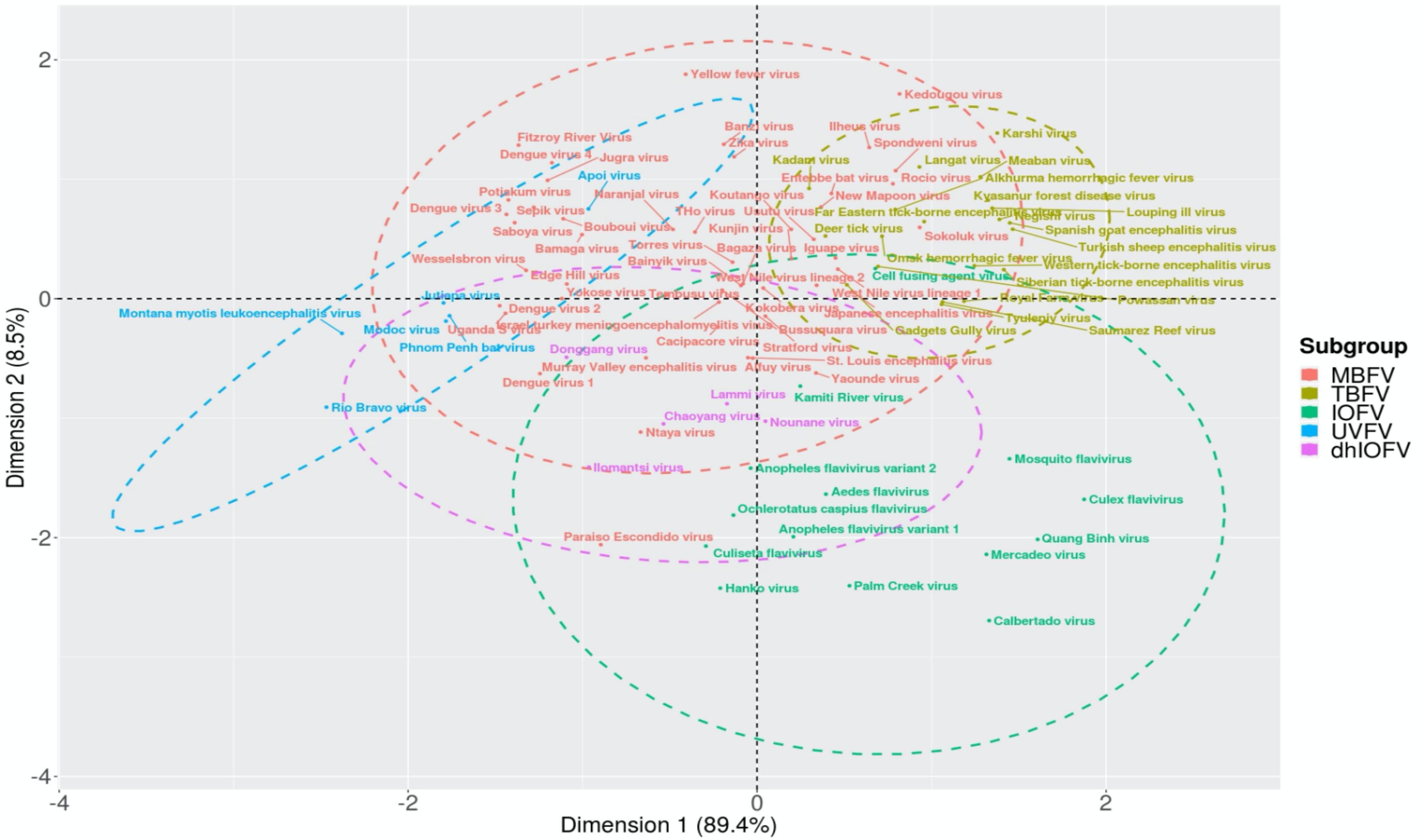


**Supplementary figure 1. Interspecies correspondence analysis and subgroup centroids of normalized Codon Adaptation Index (nCAI) values of flaviviruses, genus *Flavivirus* (N = 94).** While there is a distinct separation between the clusters of insect-only and the other flavivirus subgroups, the clouds (shown as dashed circles with colors corresponding to a flavivirus subgroup) computed from them show major overlap, indicating that most of the subgroups share similar host preferences, e.g. unknown vector flaviviruses most likely have similar hosts as mosquito-borne flaviviruses. An interesting case can be observed with Paraiso Escondido virus, which is located outside the mosquito-borne cloud, which suggests that it does not have a vertebrate host. This is supported by previous studies, in which the virus is described to infect only sandflies. MBFV = Mosquito-borne flavivirus, TBFV = Tick-borne flavivirus, IOFV = Insect-only flavivirus, UVFV = Unknown vector flavivirus, and dhIOFV = Dual-host insect-only flavivirus. The centroids were computed based on the multivariate normal distribution of each subgroup with a confidence level of 0.95. Dimension 1 explains 89.4 percent and dimension 2 contributes to 8.5 percent of the variation.

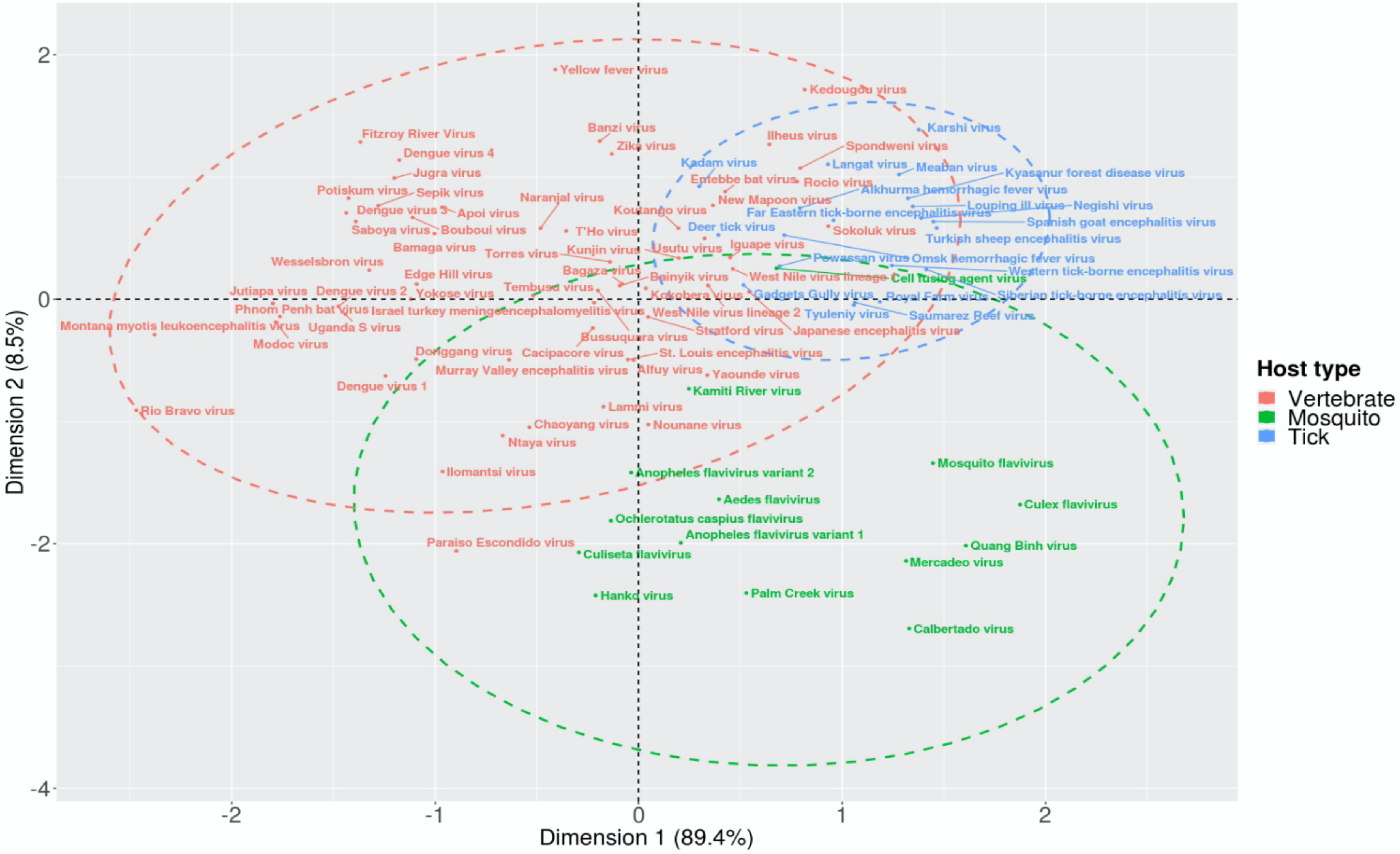


**Supplementary figure 2. Interspecies correspondence analysis with host type centroids of normalized Codon Adaptation Index (nCAI) values of flaviviruses, genus *Flavivirus* (N = 94).** When the flaviviruses are categorized based on their preferred host type, we can observe two major clouds (dashed circles) and clusters; viruses that have a vertebrate host and those that have a mosquito-host. Tick-borne viruses largely overlap with the vertebrate host cluster, suggesting that while tick-borne flaviviruses cluster separately, they have the same type of host as mosquito-borne viruses. An interesting exception in the vertebrate cluster is Paraiso Escondido virus, which is located outside the cloud. This may be due to the virus infecting mainly sandflies rather than vertebrates. The centroids were computed based on the multivariate normal distribution of each host type with a confidence level of 0.95. Dimension 1 explains 89.4 percent and dimension 2 contributes to 8.5 percent of the variation.

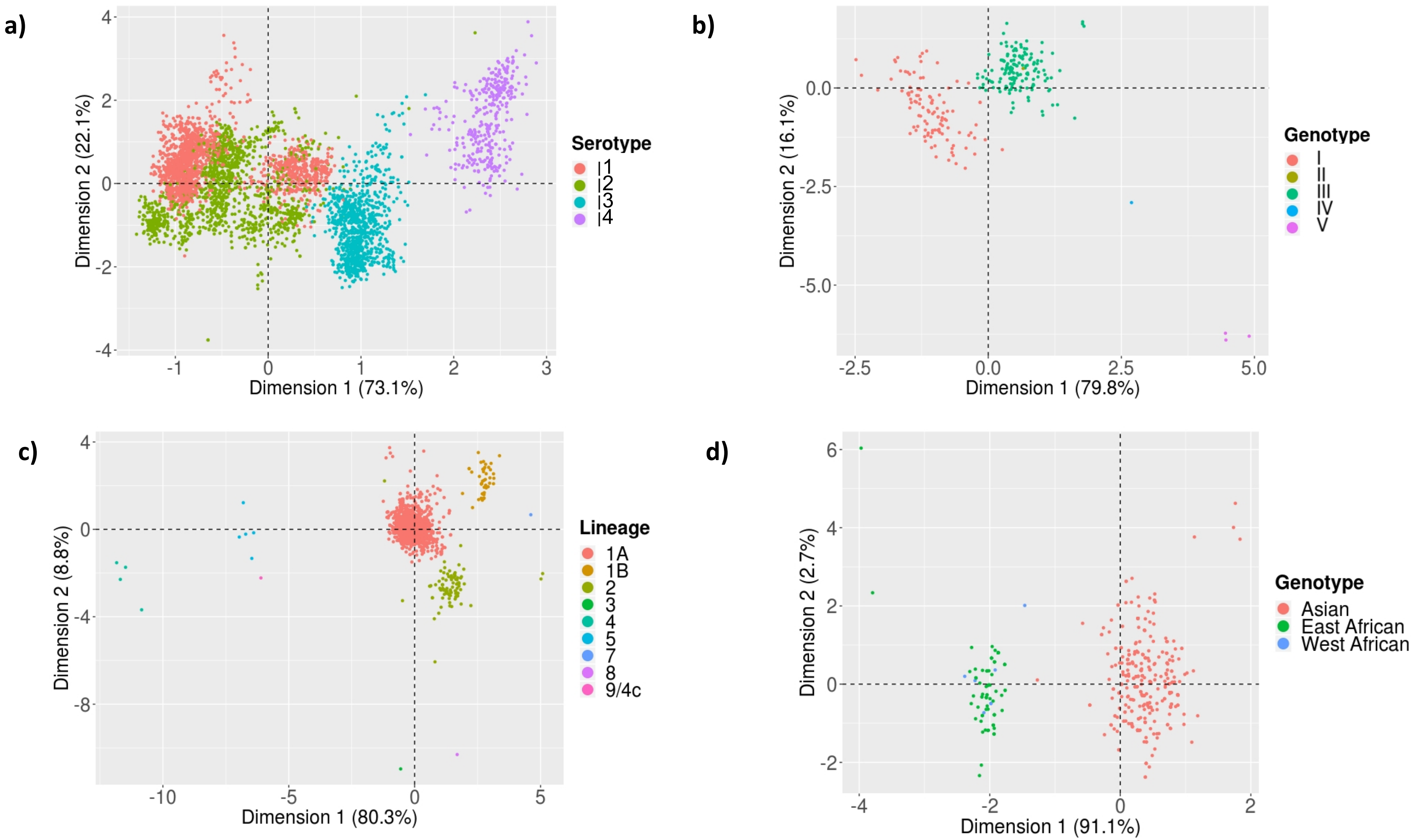


**Supplementary figure 3. Intraspecies correspondence analyses of normalized Codon Adaptation Index (nCAI) values of four major mosquito-borne flaviviruses (genus *Flavivirus*).** The results show that nCAI is able to discriminate between the different categories of (a) Dengue viruses (N = 4865), (b) Japanese encephalitis viruses (N = 297), (c) West Nile viruses (N = 1619) and Zika viruses (N = 494). Additionally, the distances between separate clusters mirror actual phylogeny. For example, Dengue virus serotypes 1–3, which are clustered close together, are more related to each other than to serotype 4, and the same can be observed with the results of the Zika virus genotypes, in which the Asian and African genotypes cluster separately, but nCAI does not distinguish East and West African genotypes from each other. In panel a, Dimension 1 explains 73.1 percent and Dimension 2 contributes to 22.1 percent of the variability. In panel b, Dimension 1 explains 79.8 percent and Dimension 2 contributes to 16.1 percent of the variability. In panel c, Dimension 1 explains 80.3 percent and Dimension 2 contributes to 8.8 percent of the variability. In panel d, Dimension 1 explains 91.1 percent and Dimension 2 contributes to 2.7 percent of the variability.

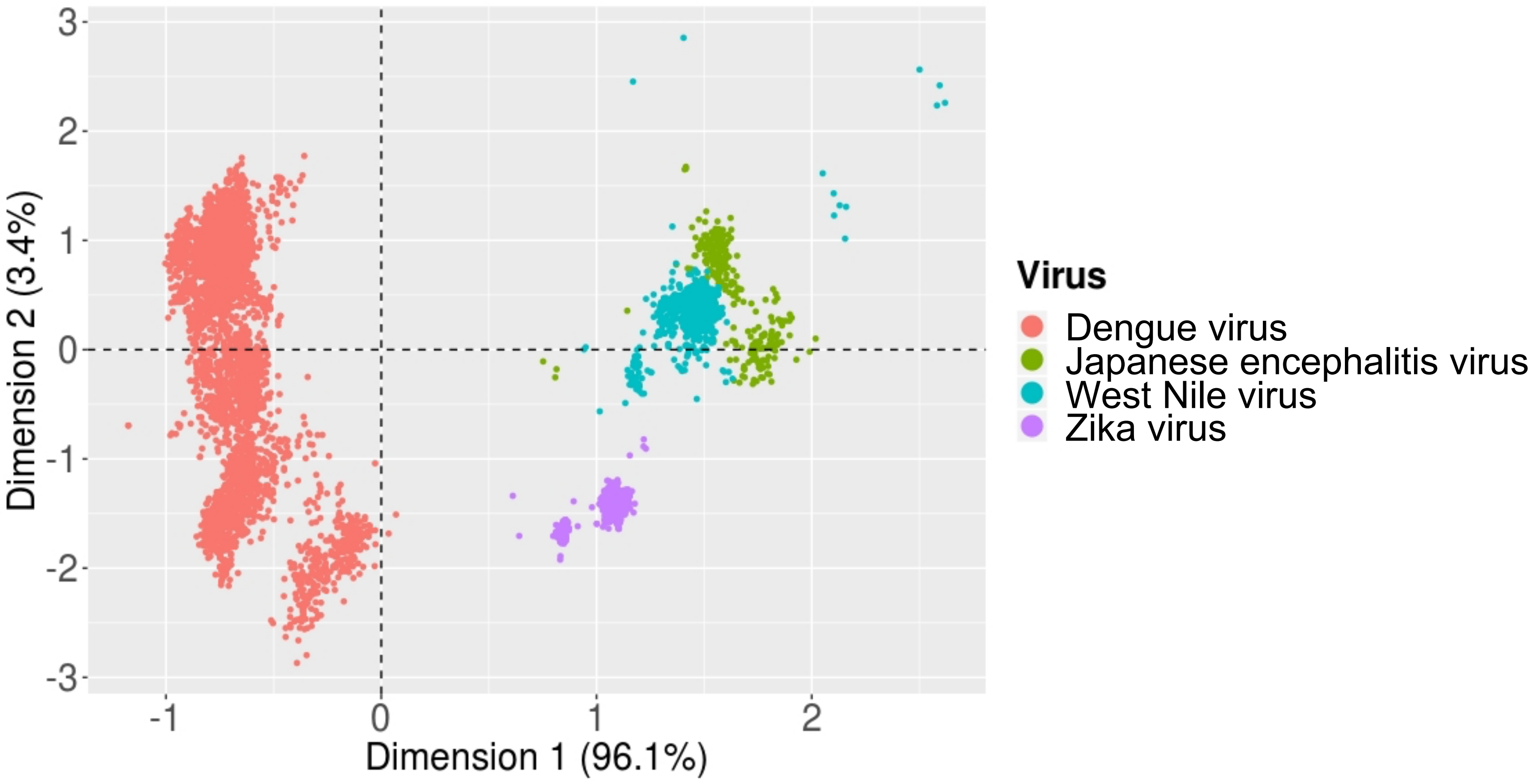


**Supplementary figure 4. Interspecies correspondence analysis of normalized Codon Adaptation Index (nCAI) values of four major mosquito-borne flaviviruses, genus *Flavivirus* (N = 7275).** The different virus genomes form distinct clusters based on computed nCAI values. The distance between these clusters mirror some phylogenetic relationships among the viruses accurately, e.g. Japanese encephalitis viruses and West Nile viruses share a more recent ancestor than to the other viruses. Of the variation, 96.1 percent is due to Dimension 1 and 3.4 percent is because of Dimension 2.

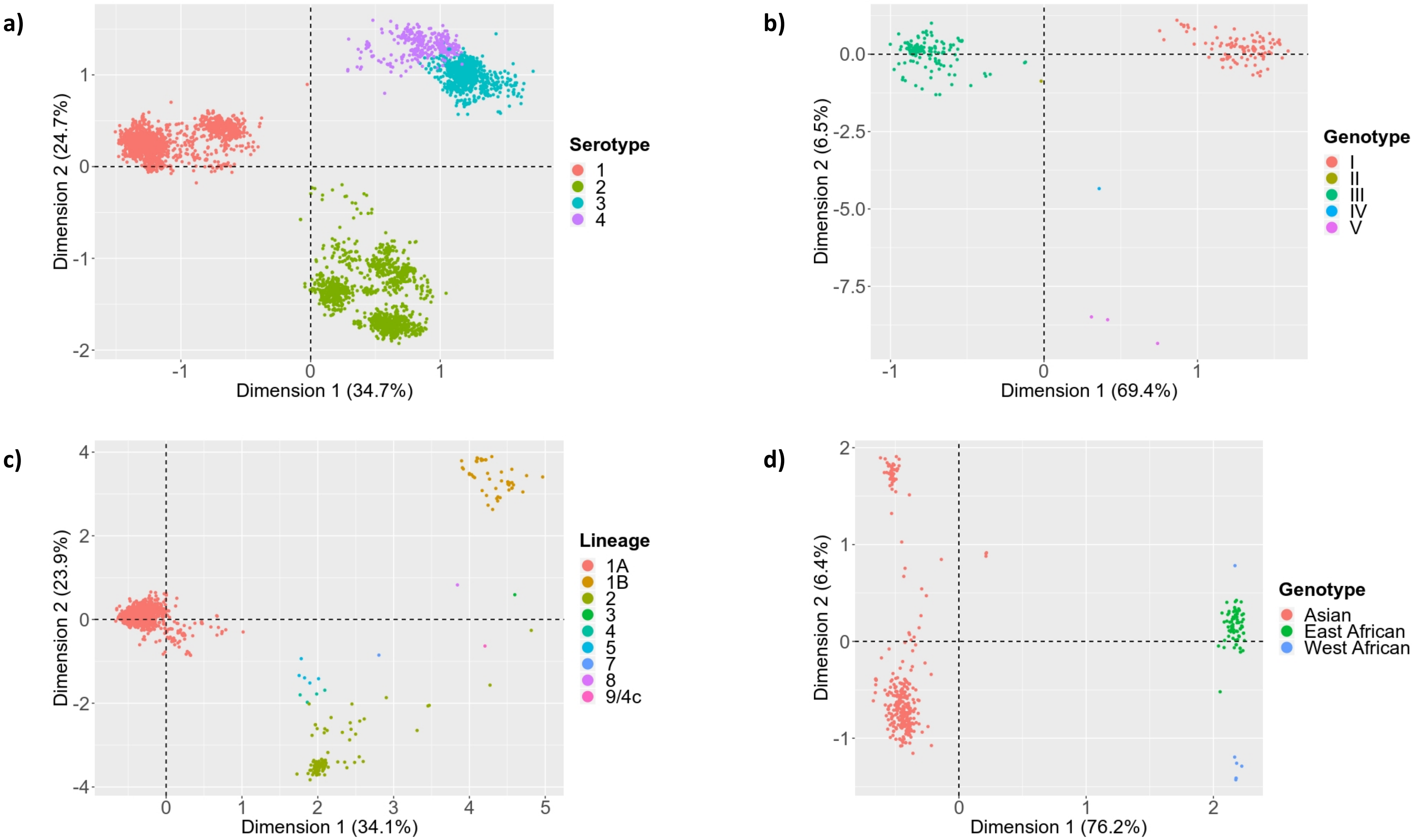


**Supplementary figure 5. Intraspecies correspondence analysis of the Relative Synonymous Codon Usage (RSCU) values of major mosquito-borne flaviviruses.** With (a) Dengue viruses (N = 4865), (b) Japanese encephalitis viruses (N = 297), (c) West Nile viruses (N = 1619) and Zika viruses (N = 494), different groups of viruses are distinguishable based on their codon usages, although the amount of separation does not necessarily mirror evolutionary relationships with the exception of Zika virus. Each virus outgroup is noted in the legends. Dimension 1 explains 34.7 percent and Dimension 2 attributes to 24.7 percent of the variation in panel a. Dimension 1 explains 69.4 percent and Dimension 2 attributes to 6.5 percent of the variation in panel b. Dimension 1 explains 34.1 percent and Dimension 2 attributes to 23.9 percent of the variation in panel c. Finally, Dimension 1 explains 76.2 percent and Dimension 2 attributes to 6.4 percent of the variation in panel d.

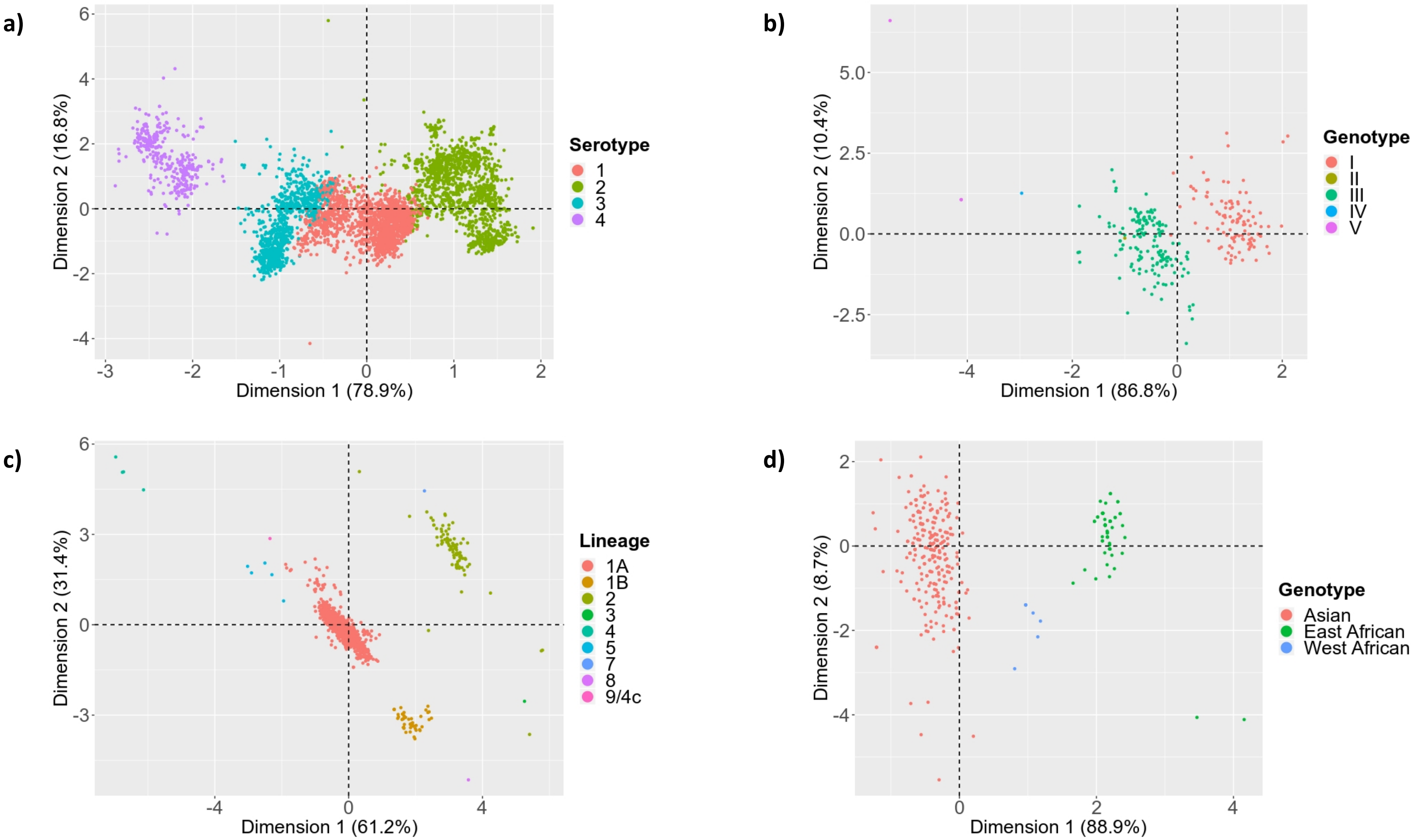


**Supplementary figure 6. Intraspecies correspondence analysis of the guanine and cytosine content based on the third nucleotide of each codon (%G3+%C3) of major mosquito-borne flaviviruses.** Based on nucleotide compositions, the genomic sequences of (a) Dengue viruses (N = 4865), (b) Japanese encephalitis viruses (N = 297), (c) West Nile viruses (N = 1619) and Zika viruses (N = 494) form distinct cluster that match established categories of each virus. Additionally, the distances between clusters is similar to their phylogenetic relatedness. Each outgroup virus of a respective flavivirus is mentioned in the legend. In panel a, 78.9 percent of the variation is explained by Dimension 1 and 16.8 percent by Dimension 2. In panel b, 86.8 percent of the variation is explained by Dimension 1 and 10.4 percent by Dimension 2. In panel c, 61.2 percent of the variation is explained by Dimension 1 and 31.4 percent by Dimension 2. In panel d, 88.9 percent of the variation is explained by Dimension 1 and 8.7 percent by Dimension 2.

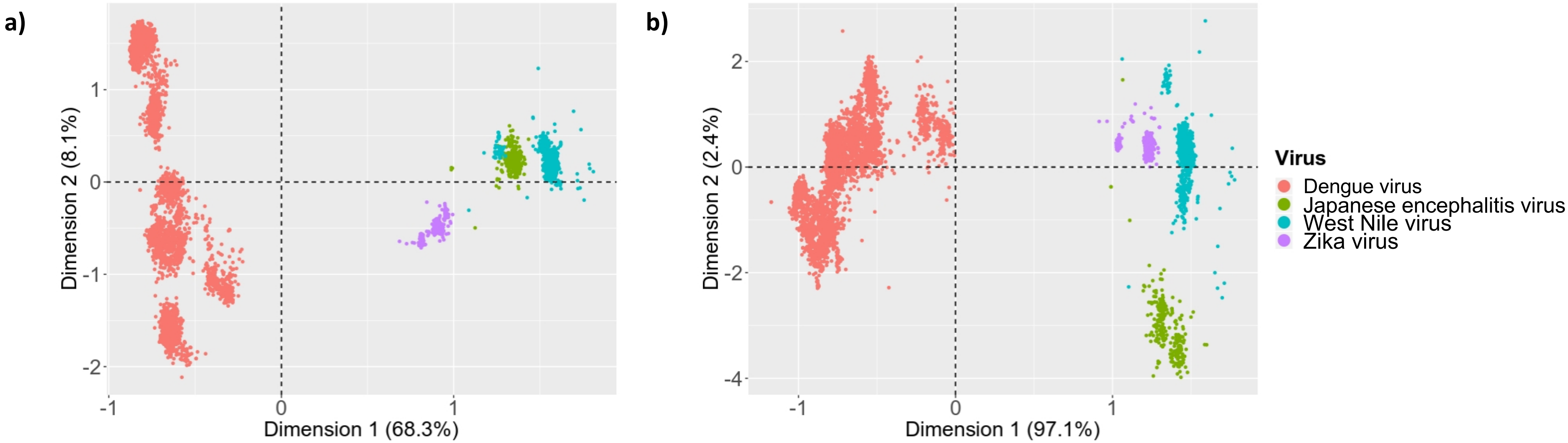


**Supplementary figure 7. Interspecies correspondence analysis of the genomic compositions of major mosquito-borne flaviviruses (N = 7275).** The viruses form species specific clusters based on their (a) Relative Synonymous Codon Usage (RSCU) values and (b) guanine and cytosine contents based on the third nucleotide of each codon (%G3+%C3). While both methods are capable of distinguishing subgroups within a species, RSCU has a greater ability to separate these groups compared to %G3+%C3. The distances between clusters do not perfectly equate to phylogenetic relationships, e.g. in panel b, Zika viruses are not more related to West Nile viruses than to Dengue viruses. With both analyses, Cell fusing agent virus was used as an outgroup. In panel a, Dimension 1 and 2 explain 68.3 and 8.1 percent of the variation respectively. In panel b, Dimension 1 contributes to 97.1 percent of the variance, while Dimension 2 contributes to 2.4 percent.

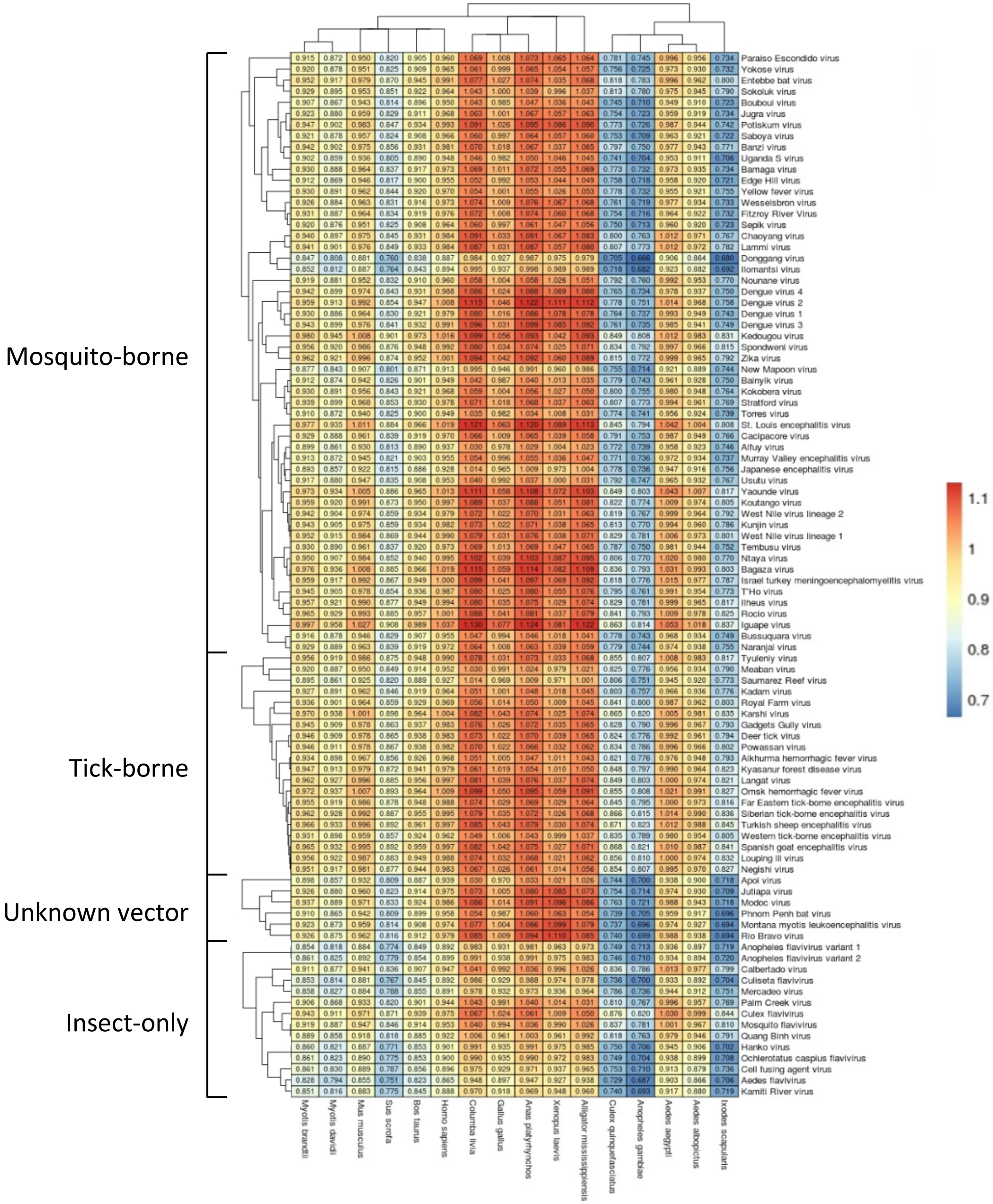


**Supplementary figure 8. Normalized Codon Adaptation Index (nCAI) heatmap of flavivirus (genus *Flavivirus*) subgroups (N = 94).** The nCAI values show overall over-optimization (nCAI > 1.05) for avians, reptiles and amphibians, and under-optimization for *Culex* and *Anopheles* mosquitoes (nCAI < 0.95). Optimal hosts tend to be mammals and *Aedes* mosquitoes (nCAI 0.95–1.05). The columns are sorted according to taxonomic classification of hosts, and the rows are in accordance with the phylogeny of flaviviruses.

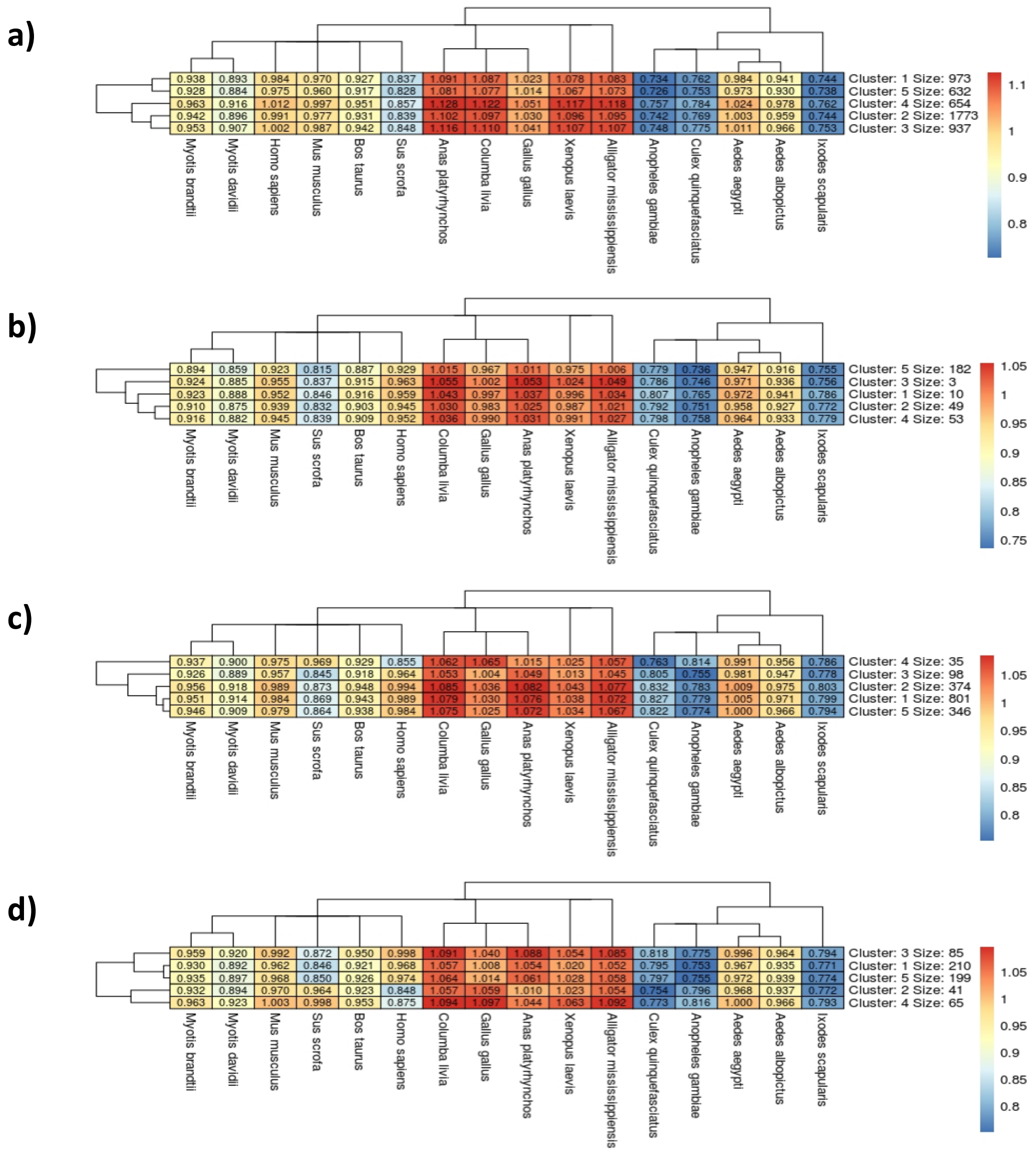


**Supplementary figure 9. Intraspecies k-means (5) heatmaps of normalized Codon Adaptation Index (nCAI) values of major mosquito-borne flaviviruses (genus *Flavivirus*).** When the nCAI values of (a) Dengue viruses, (b) Japanese encephalitis viruses, (c) West Nile viruses and (d) Zika viruses are plotted in heatmaps, they display similar overall adaptation levels to different host organisms, although there are differences between these viruses. On average the viruses are optimized for mice *(Mus musculus)* and humans *(Homo sapiens),* and *Aedes* mosquitoes, especially *Aedes aegypti* (nCAI 0.95–1.05), thus being likely hosts. The unlikely hosts are the other mammalian hosts, avians, reptiles and amphibians due to over-optimization (nCAI > 1.05), and *Culex* and *Anopheles* mosquitoes, and ticks *(Ixodes scapularis)* due to under-optimization (nCAI < 0.95). There are however exceptions; Japanese encephalitis viruses in panel b have as more likely optimal hosts mostly birds, reptiles and amphibians. The number and size of k-means clusters do not match the current classification of these flaviviruses.
